# Massively parallel encapsulation of single cells with structured microparticles and secretion-based flow sorting

**DOI:** 10.1101/2020.03.09.984245

**Authors:** Joseph de Rutte, Robert Dimatteo, Maani M Archang, Mark van Zee, Doyeon Koo, Sohyung Lee, Allison C. Sharrow, Patrick J. Krohl, Michael P. Mellody, Sheldon Zhu, James Eichenbaum, Monika Kizerwetter, Shreya Udani, Kyung Ha, Andrea L. Bertozzi, Jamie B. Spangler, Robert Damoiseaux, Dino Di Carlo

## Abstract

Techniques to analyze and sort single cells based on functional outputs, such as secreted products, have the potential to transform our understanding of cellular biology, as well as accelerate the development of next generation cell and antibody therapies. However, secreted molecules rapidly diffuse away from cells, and analysis of these products requires specialized equipment and expertise to compartmentalize individual cells and capture their secretions. Herein we demonstrate the use of suspendable microcontainers to sort single viable cells based on their secreted products at high-throughput using only commonly accessible laboratory infrastructure. Our microparticles act as solid supports which facilitate cell attachment, partition uniform aqueous compartments, and capture secreted proteins. Using this platform, we demonstrate high-throughput screening of stably- and transiently-transfected producer cells based on relative IgG production as well as screening of B lymphocytes and hybridomas based on antigen-specific antibody production using commercially available flow sorters. Leveraging the high-speed sorting capabilities of standard sorters, we sorted >1,000,000 events in less than an hour. The reported microparticles can be easily stored, and distributed as a consumable reagent amongst researchers, democratizing access to high-throughput functional cell screening.

## Introduction

The microwell plate is a foundational component of biological assays because of its ability to scale experiments and integrate with lab automation infrastructure. This simple piece of plasticware allows scientists to add or exchange reagents while preventing cross-talk between samples. The bottom surface of each well can be functionalized to adhere cells, promote cell growth, and perform biomolecular reactions that allow for colorimetric and fluorescent readouts. Biological samples of interest can then be isolated by pipetting fluid from a well. Despite its simplicity and utility, even the highest throughput, 1536-well plate formats, hold volumes that are hundreds of thousands of times larger than individual cells, limiting sensitivity and throughput in studies of functional properties, such as secreted products, from single cells.

The ability to perform functional biological assays at single-cell resolution promises to deepen our understanding of biology and accelerate the development of new biotechnology products. Among the ∼20,000 protein-coding genes in the human genome more than 15% of the encoded proteins are predicted to be secreted, a similar percentage to that for cell membrane-bound proteins (*1*). New approaches to analyze and sort cells based on these secreted factors can shine light on this ubiquitous but significantly understudied cellular function. In biotechnology, sorting of rare B cells or plasma cells directly based on the secretion of antigen-specific antibodies allows for the acquisition of gene sequences that can be used to make new monoclonal antibody (mAb) drugs or diagnostic affinity reagents (*2*–*7*). Additionally, selection of stable production-grade cells such as Chinese hamster ovary (CHO) cells based on humanized IgG secretion rates enables development of highly productive cell lines for industrial scale monoclonal antibody manufacturing (*8*).

Although technologies have emerged that enable the interrogation of single cells, there are significant tradeoffs that either limit their functionality or inhibit widespread adoption. Fluorescence activated cell sorting (FACS) was one of the first single-cell technologies to gain widespread adoption, enabling users to probe and sort individual cells based on scattered light and fluorescently tagged molecular labels at throughputs of over 10,000 events per second. Screening of viable cells is typically limited to cell surface markers (i.e. the clusters of differentiation markers, or CDs) as analysis of intracellular markers requires membrane permeabilization and fixation (*9, 10*). Approaches have been developed to sequester secreted products directly onto the cell surface, but these techniques suffer from significant crosstalk between cells (*11*–*13*). Microfluidic technologies have emerged that can create volumes approaching the size of cells, enabling high-throughput analysis of single cells based on additional properties such as secreted molecules (*14*). Devices with microscale chambers as well as water-in-oil droplets generated using microfluidic devices have been employed to isolate cells, accumulate secreted molecules, and even sort (*15*–*20*). Despite the utility of these existing platforms, the ability to add reagents and wash is limited, and microfluidically-generated droplets lack the solid surface of a microwell plate which is critical for standard assay formats like sandwich immunoassays or ELISA. Strategies to address these shortcomings have been developed, but each requires specialized assay formats, and often extremely expensive equipment to perform the assays, which hinders widespread adoption (*21*–*24*).

We introduce a facile approach to perform functional assays on individual cells in high throughput using structured microparticles, which act as suspendable and sortable microwells (Figure 1). These microparticles, or “nanovials”, hold single cells in sub-nanoliter volumes of fluid, 100,000 times less volume than a single well of a 1536-well microwell plate, yet require no specialized instrumentation. Fluids are easily exchanged by centrifugation and pipetting, and each compartment can be sealed and unsealed using biocompatible oils to prevent cross-talk between samples. The surfaces can be modified to bind cells or capture biomolecules for various molecular readouts. Nanovials can be analyzed and isolated using commonly available FACS instruments enabling screening at rates >1000 events per second. While other particle systems have been utilized to hold sub-nanoliter volumes (*25*–*29*), ours is the first approach that allows attachment and protection of cells in cavities within particles, unlocking new capabilities to expand the scale of analysis by integration with flow cytometry. Using this easily-adopted nanovial format we conducted a screen of over 1 million events in less than an hour using a commercial FACS instrument, sorting out rare antibody-secreting cells from orders of magnitude more abundant background cells, all within one day. After sorting, cells remain intact and viable, enabling regrowth and single-cell RT-PCR. A similar experiment would require >2600 384-well plates and several weeks of additional culture to grow up a sufficiently large clonal population of cells required to detect secretions in the larger volumes. Nanovial-enabled workflows promise to democratize antibody discovery and cell line development, and also empower researchers to investigate single-cell secretions, as a key cellular function, with unprecedented precision and scale.

**Figure 1.**
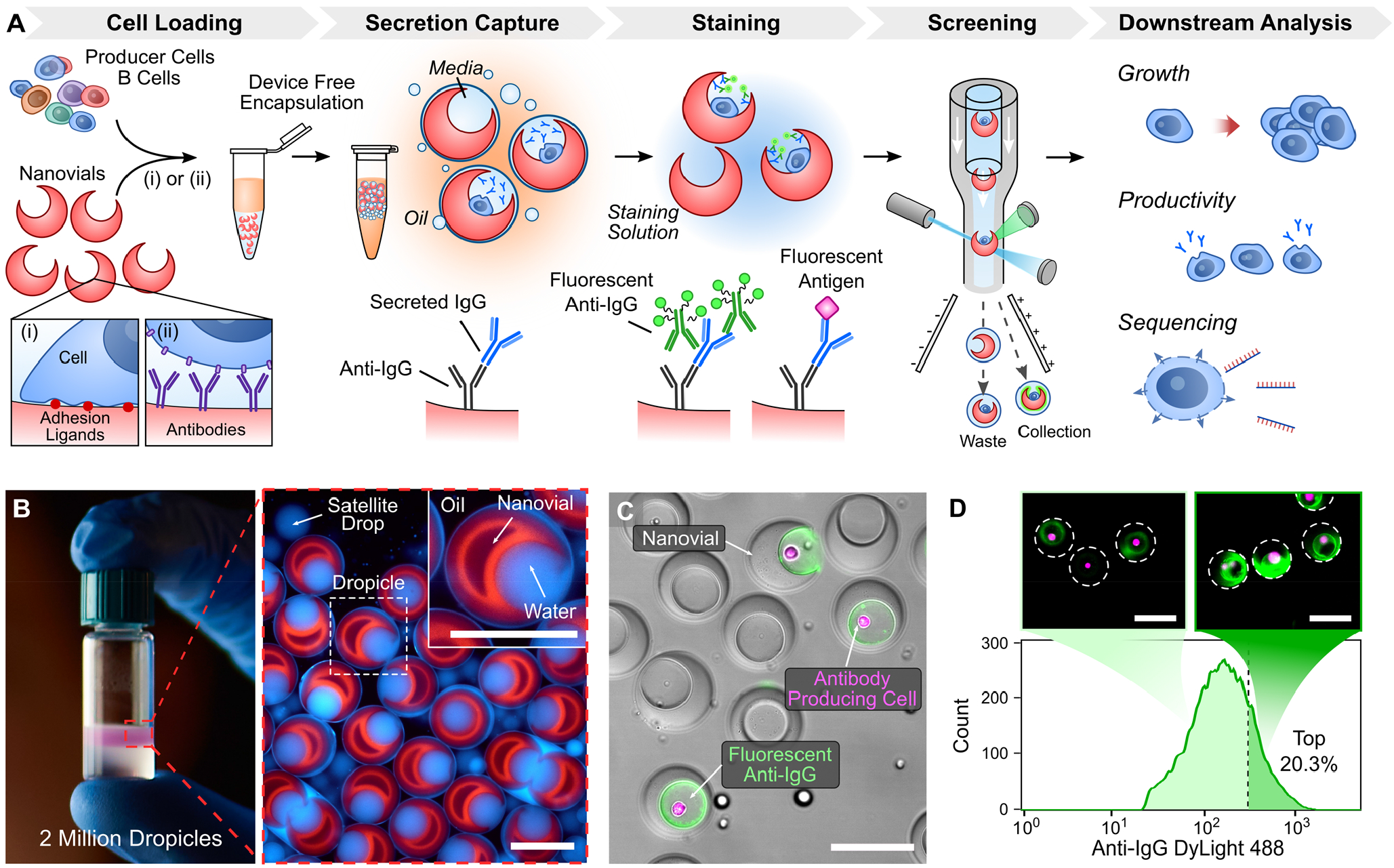
Microparticle platform for high-throughput single-cell secretion screening. **(A)** Individual cells are loaded into prefabricated microparticle containers (nanovials) and bound using a variety of binding schemes (i,ii). Particles and associated cells are agitated by pipetting with biocompatible oil and surfactant to generate monodisperse compartments (known as dropicles), which are dictated by the particle size preventing crosstalk between microcontainers. Cells are incubated to accumulate secretions on associated particles and transferred back to a water phase for fluorescent labeling. Particles, cells, and associated secretions are then analyzed and sorted using high-throughput commercial flow sorters. Isolated populations can then be screened downstream for different phenotypic (growth, productivity) and genotypic properties (RNA expression). **(B)** Photograph of a tube with 2 million dropicles formed using simple pipetting steps in less than 1 minute (left). Fluorescence microscopy image of uniform dropicles formed with fluorescently-stained nanovials and water-soluble dye (right) show distinctly sealed sub-nanoliter volumes. **(C)** Microscopy image of nanovials with cells and associated secretions after unsealing the nanovials and staining with fluorescent labels. **(D)** Example of a flow cytometry plot and post-sort images of enriched highly-secreting populations. Scale bars are 100 µm.

## Results

### Precise fabrication of suspendable microcontainers

Our approach to screen single cells requires cavity-containing hydrogel microparticles that are engineered to directly load and protect cells. Each particle is decorated with functional groups to attach cells and perform chemical reactions, is precisely shaped to prevent cross-talk, and is designed to be compatible with commercial flow cytometers. Utilizing an aqueous two-phase system combined with droplet microfluidics, and hydrogel chemistry developed for tissue engineering, we fabricate these hydrogel microparticles (nanovials) at high throughputs (∼1000 s^-1^) (Figure S1-3) (*30, 31*). By tuning the fabrication parameters, we achieve highly monodisperse polyethylene glycol (PEG)-based nanovials with accessible internal cavities (outer diameter CV of 1.5 - 3.2%, cavity opening diameter CV of 2.1 – 5.8%, depending on size Figure S1F-G). Particles are easily modified with various cell adhesion moieties (e.g. argine-glycine-aspartate (RGD), poly-L-lysine (PLL), fibronectin). Biotin covalently coupled to the nanovial matrix allows for facile surface decoration with streptavidin (Figure S1G) which allows linkage with biotinylated antibodies or antigens that bind to secreted products. High-affinity biotin-streptavidin interactions can also be used to link cell capture antibodies specific to cell surface proteins (e.g. CD45 and CD19) to selectively adhere cells to nanovials. Biotin-functionalized cells can also be directly captured on streptavidin-coated nanovials. The morphology of the nanovial cavity and relative size can be adjusted by tuning the concentration of components in the particle precursor solutions (Figure S2). We are able to fabricate monodisperse nanovials across a broad range of mean diameters (Figure S3) from 35 to 83 µm, which are compatible with a range of cell types and common lab and FACS instruments (Figure S2). Importantly, this nanovial fabrication step is the only part of the process that requires use of a specialized microfluidic device, and fabrication is completely decoupled from the biological assays themselves. All parts of the single-cell assay workflows are performed directly with pre-fabricated nanovials and standard lab equipment (Video S1). This is a critical feature of our platform as it enables the more complex microfluidic/particle fabrication work to be centralized and performed in advance.

### Nanovials as modular single-cell carriers

Loading cells into the cavities of the nanovials is achieved with simple pipetting steps followed by incubation to allow cell binding. Nanovials can be first loaded into a standard well plate by pipetting. Due to their unique morphology, nanovials settle with their cavities mostly upright, forming a monolayer of particles, with exposed cavities, at the bottom of the well (Figure S1H, S4, Video S2) (*32*). Cells are then seeded over the particles and settle with a sizeable fraction coming to rest in the particle cavities (Figure 2). Notably, this seeding approach led to cell occupancies that closely followed Poisson statistics (Figure 2D), as is expected for loading of single cells into microfluidically-generated droplets or for limiting dilution loading of single cells into microwells (*33, 34*). By controlling the cell seeding density, we may control occupancy such that most particles have either 0 or 1 cells associated with them.

**Figure 2.**
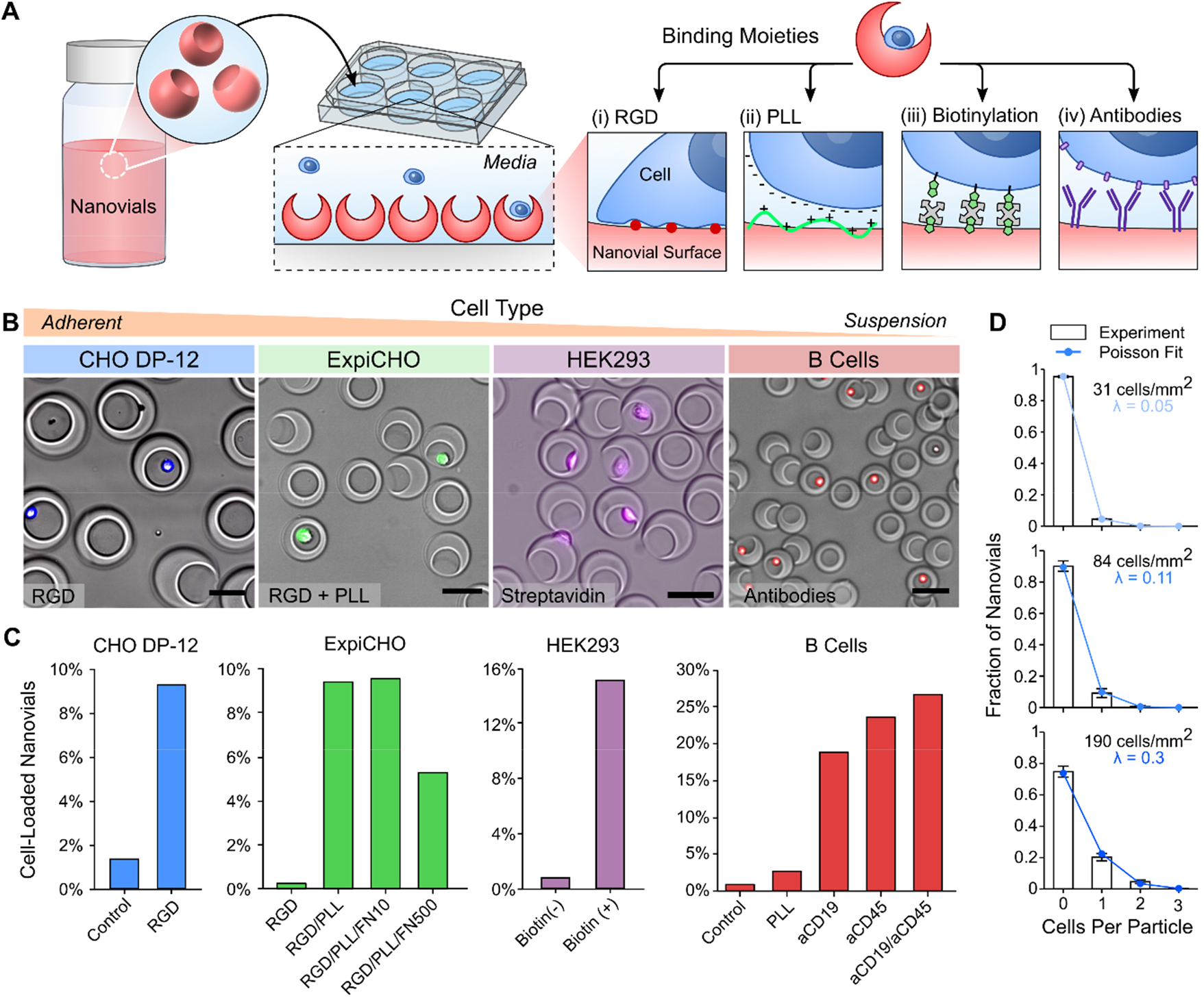
Loading and binding of single cells into nanovials. **(A)** Particles are loaded into wells and settle with their cavities oriented upright due to their asymmetric center of mass. Cells are then seeded into the open cavities and attach to various binding moieties. Unbound cells are washed away using a reversible cell strainer and recovered particles and associated cells are analyzed. **(B)** Example microscopy images of fluorescently tagged cells bound to nanovials. **(C)** Comparison of cell binding is plotted below for different cell types and binding moieties. Adherent CHO DP-12 (blue) cells are bound to particle cavities through integrin binding sites (RGD peptide) linked to the particle matrix. Cell retention for suspension adapted ExpiCHO (green) was increased by introduction of positively charged poly-L-lysine (PLL). Biotinylated HEK293 cells bound to streptavidin coated nanovials (purple). Inclusion of surface-marker-specific antibodies on the nanovial surface increased retention of B cells (red). **(D)** Loading of cells into nanovial cavities follows Poisson statistics. > 500 particles were analyzed for each condition in cell retention experiments. Error bars represent SD of n = 3 samples for loading distribution.

Following seeding, cells can be bound to the nanovials through one of a variety of cell-surface interactions, that are tailored to the cell type of interest. Adherent producer cell lines used for the production of mAbs and other biologics can be adhered through cell surface interactions with RGD, a well-known adhesive peptide motif present in fibronectin. CHO DP-12 and human embryonic kidney (HEK) 293 cells adhered (Figure 2B,C) and maintained high levels of single cells within cavities following vigorous wash steps. Suspension-adapted cell lines, such as ExpiCHO cells, did not adhere well to RGD alone. However, increased adhesion similar to other cell lines was achieved by modifying particles to also contain poly-L-lysine (PLL) (Figure 2B). For other primary suspension cell populations with well-defined surface market expression, such as B cells, we found that antibodies against cell surface proteins led to optimum adhesion in nanovial cavities even after vigorous washing and sorting. Antibodies against CD45, CD19, or both yielded high levels of B cell adhesion and maintenance that followed Poisson loading statistics (Figure 2C). We also showed that surface proteins of HEK293 cells can be biotinylated such that the cells adhered and were maintained on nanovials through biotin-streptavidin interactions (Figure 2B,C).

### Sealing of nanovials using biocompatible oil and surfactants to prevent cross-talk

Uniform droplets can be formed around nanovial particles via simple pipetting steps with oil and surfactant to seal them and prevent cross-talk between samples (Figure 3). To accomplish this, an aqueous solution of nanovials is first concentrated in a conical or microcentrifuge tube by centrifuging and aspirating the supernatant. A layer of biocompatible oil with surfactant is added and the suspension is then pipetted vigorously for 1 min to create smaller and smaller water-in-oil droplets. Eventually the droplet size is maintained by the outer periphery of the microparticle and we find that, in agreement with minimal energy considerations (Figure S5) (*35*), uniform volumes of fluid remain stably trapped within the cavity by the hydrophilic particle. Any excess fluid in the suspension is partitioned into much smaller satellite droplets (Figure 3B). The resulting emulsion shows two unique distributions: (i) small non-uniform satellite droplets and (ii) monodisperse droplets each templated by single particles (Dropicles) (Figure 3C). This compartmentalization process was tested successfully over a range of nanovial sizes (35-85 µm diameters). For larger particles, it was found that uniform emulsions (mean diameter of 99 µm, CV = 4%) could be formed directly from concentrated nanovials with minimal aggregates (<1%, Figure 3D). However, for smaller nanovials an increase in particle number density in the pelleted sample led to numerous droplets containing multiple particles. We found that dilution of the sample to the same total number of particles per unit volume remedied this particle aggregation, enabling the formation of smaller uniform partitions (mean diameter of 40 µm, CV = 3.2%, <5% aggregates, Figure 3).

**Figure 3.**
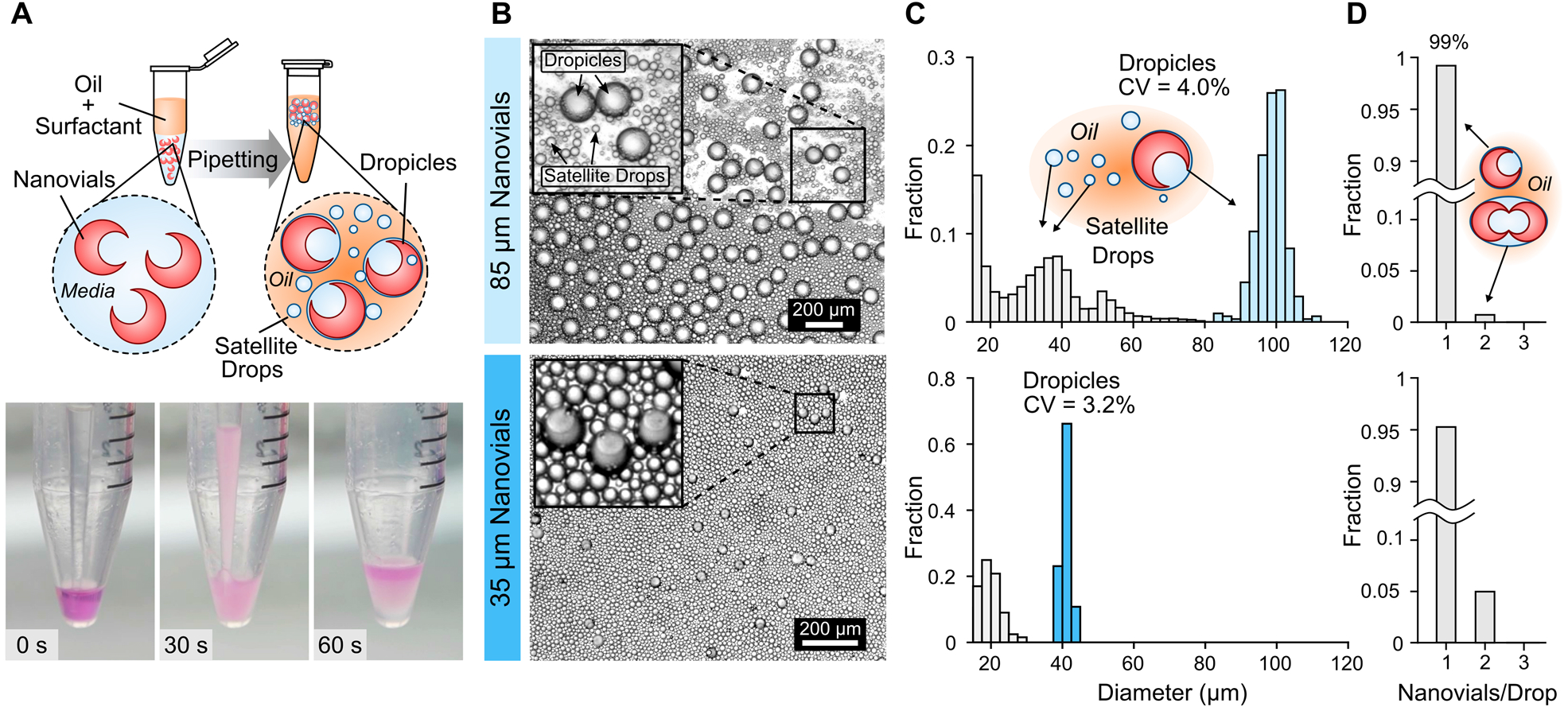
Massively parallel device-free formation of uniform droplets. **(A)** Biocompatible oil and surfactant are added to a tube containing concentrated particles. The suspension is pipetted vigorously for 1 minute to generate smaller and smaller emulsions. **(B)** Microscopy images of the emulsions show a uniform population of droplets containing particles and smaller background satellite droplets. **(C)** Histograms of dropicle diameter for different nanovial sizes. Nanovial-containing droplets are highly uniform with comparable size distributions to advanced microfluidic techniques (CV<5%). **(D)** For both nanovial sizes, nearly all droplets formed have either 0 or 1 particle encapsulated, with only a small fraction (<1% for 85 µm particles and <5% for 35 µm particles) containing 2 or more particles per droplet. Scale bars, 200 µm.

Importantly, nanovials can be recovered from the oil while still retaining functional cells. To recover nanovials a biocompatible destabilizing agent is used to coalesce the droplets together (Figure S6C). Live/dead analysis of CHO cells following encapsulation and subsequent emulsion breaking showed high viability over a 24-hour period (>80%) indicating that the workflow is biocompatible (Figure S6A-B). Further, we observed similar growth rates from cells expanded after dropicle release in comparison to cells seeded directly into a well plate (Figure S6C-F). CHO DP-12 cells adhered via integrin binding and B cells bound via antibodies remained attached to the nanovial cavities over the entire emulsification process. After washing and emulsification, exceedingly few cells are adhered to the external surface of the nanovials suggesting these cells are sheared off and that the cavity can shelter cells from shear forces during processing.

### Single cell secretion analysis and sorting using nanovials

Using the nanovial platform, we demonstrate a device-free workflow to perform single-cell secretion assays with minimal crosstalk (Figure S7). Given the importance of selecting cell lines with high antibody titer for therapeutic production(*36*), we chose a model system comprised of a Chinese hamster ovary (CHO) cell line that produces human IgG targeting interleukin-8 (IL-8) and particles modified to capture human IgG. Cells are first loaded into the microparticle cavities and adhere via integrin binding sites as previously described. After initial cell seeding, particles and associated cells are collected and washed to remove background secretions. Particles are then coated with anti-human IgG Fc antibodies by binding to biotin groups linked to the particle matrix. Following this step, the nanovials and associated cells are rapidly compartmentalized in less than 1 minute by pipetting with oil and surfactants. The compartmentalized CHO cells are then incubated, and the secreted antibodies are captured onto the associated particle matrix. After incubation, the emulsions are broken and the nanovials containing attached cells and secretions are collected and washed. Nanovials are then labeled with secondary fluorescent antibodies targeting the secreted anti-IL-8 antibodies. Using fluorescence microscopy, we confirmed retention of cells and associated secretion signal on the particle surface after recovering out of oil into an aqueous phase (Figure 1C).

Because of the high fraction of secreting cells, preventing secretion cross-talk between cells in neighboring particles is critical to enable quantitative analysis and sorting based on secretion differences at the single-cell level. Rapid encapsulation is key to preventing cross-talk of secretions, as demonstrated by a side-to-side comparison of the secretion assay with and without the oil encapsulation step (Figure 4A-B). When secretions are captured on particles without the encapsulation step, particles without associated cells show high secretion signals indicating significant cross-talk (8.2% of empty particles have signal above threshold) (Figure 4A). Conversely, with the oil encapsulation step there are two visibly distinguishable populations, nanovials with and without secreting cells, and only 1% of nanovials without cells have signal above the cutoff threshold (Figure 4B). Minimizing the fraction of false positives is particularly critical when secretions can spill over into neighboring nanovials containing non-secreting cells that would contaminate downstream cultures or sequencing assays.

**Figure 4.**
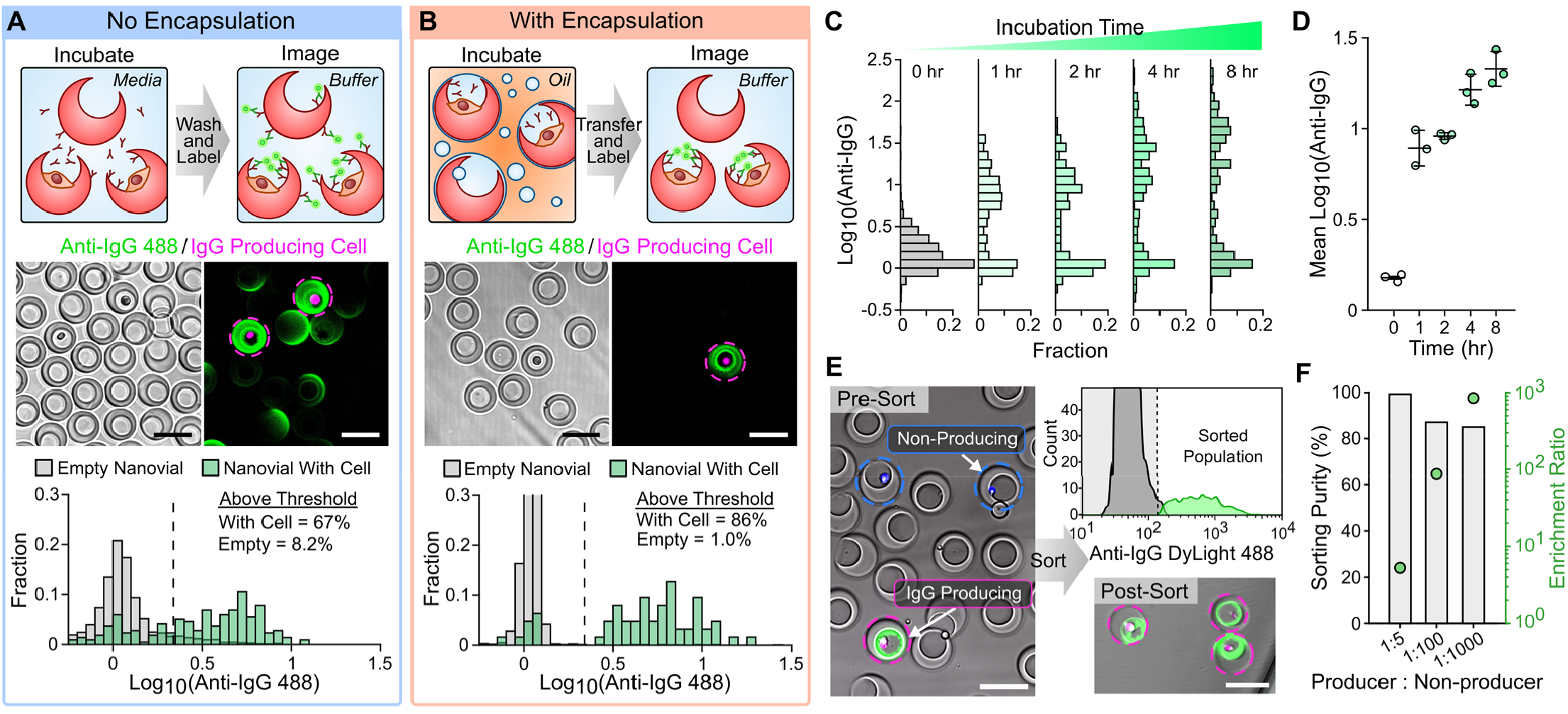
Analysis of single-cell secretions using dropicles. **(A-B)** Characterization of cross-talk when performing a secretion assay with human IgG producing CHO cells with and without an encapsulation step. **(A)** Significant cross-talk is observed when cells and particles are left in an aqueous phase during secretion incubation. **(B)** Minimal cross-talk is observed when cells and associated particles are encapsulated in oil during the secretion incubation step. Thresholds in both **(A)** and **(B)** are set at 3 standard deviations above empty particle signal for the encapsulation condition. **(C)** Measured secretion signal increases with incubation time due to accumulation of more secreted molecules. **(D)** Averaged secretion measurements across the producing population shows signal increasing proportionally with time across 3 separate samples. Error bars represent standard deviation between samples. **(E)** A secretion assay was performed on a mixture of human IgG producing CHO cells (magenta) spiked into a non-producing CHO cell population (blue). Samples were sorted based on positive IgG signal using FACS and imaged using fluorescence microscopy to determine purity and enrichment ratio. **(F)** Spiked target cells were successfully isolated across a range of dilutions (1:5 – 1:1000) at purity up to 99% and nearly 1000-fold enrichment. 100,000 single cells were sorted during the enrichment study.

We observe an increase in mean accumulated fluorescence signal on the particles that trends, as expected, with increasing incubation time and corresponding antibody production (Figure 4C-D). Using fluorescence microscopy, we detected significant signal over background (3 standard deviations) after 1 hour of incubation. Interestingly, we noted across all incubation times that there was a sizeable population of cells (∼25%) that had no measurable secretion signal (Figure 4C). Viability analysis of this population revealed that >50% of these cells were still alive indicating that the lack of signal is likely due to a fraction of the cell population no longer expressing the antibody production gene or secreting the produced antibodies.

In addition to characterizing IgG production for the stable CHO cell line, we also demonstrated the ability to characterize the distribution of secreted product from transiently transfected HEK293 cells (Figure S8A). HEK293 cells were transfected with plasmids coding for two recombinant IgGs: atezolizumab and 10H2. Cells were first loaded onto streptavidin-coated nanovials via surface biotinylation of HEK293 cells, and plasmid was introduced. Cells and associated nanovials were then emulsified and cells were allowed to secrete recombinant products for 32 hrs. The emulsion was subsequently broken and nanovials were recovered for flow cytometry (Figure S8B-C). For atezolizumab and 10H2, 24.2% and 44.7% of cell-containing nanovials, respectively, showed antibody signal above background empty nanovials (Figure S8D), demonstrating minimal crosstalk in the system. We also observed a long tail of secretion profiles for both atezolizumab and 10H2, suggesting substantial heterogeneity in the level of gene expression and/or functional secretion. Similarly, 38.1% and 27.6% of cell-containing nanovials for the atezolizumab and 10H2 transfected conditions, respectively, showed antibody signal above threshold defined by the non-transfected control (Figure S8E).

To further validate the capacity of our system to identify rare subpopulations based on secretions, a FACS enrichment experiment was performed. We performed secretion-based sorting with a mixed population of the anti-IL-8 producing CHO cells and non-producing CHO cells each labeled with a separate Cell Tracker™ dye (Figure 4E). After the secretion assay and prior to sorting, fluorescence imaging of the stained 85 µm diameter particles showed increased signal on particles containing the antibody-secreting cells of interest compared to those containing non-secreting cells (Figure 4E). This further demonstrated the lack of cross-talk in our system as well as the specificity of the labels to the secretions of interest. The particles with associated cells and secretions were then sorted based on the labeled secretion intensity (Figure 4E). Downstream analysis showed effective isolation of the sub-population of interest with high-purity (85-99%) (Figure 4F) over a range of target cell dilutions. In the most dilute case (1:1000) an enrichment ratio of 850-fold was achieved indicating the capability to isolate rare target cell events at frequencies of ∼0.1%. Cells isolated using this approach can be expanded directly from the particle matrix enabling a streamlined workflow with minimization of trypsinization steps (Figure S6C-D). Collectively, our stable and transient transfection studies illustrate the versatility of the nanovial technology for identifying and recovering rare populations of high-yielding producer cells using standard flow cytometry workflows.

### Viable enrichment of high-titer subpopulations using FACS

We hypothesized that sorted cells would maintain the ability to produce antibodies at levels correlating with the mean intensity of initial single-cell secreted signal. To test this hypothesis, we performed the nanovial secretion assay on human anti-IL8 producing CHO cells and selected out sub-populations based on level of IgG secretion signal using FACS (Figure 5). Cells secreting antibodies were sorted in high-throughput (>200 events/s) by gating off both the fluorescently labeled secretion channel as well as CellTracker™ dye (Figure 5A). For each separate passage of cells analyzed (n = 4) we sorted two sub-populations: (1) all particles with cells and detectable secretion signal above background and (2) particles with cells and the top 20% of antibody secretion signal. Microscopy images of the sorted nanovials and associated cells show successful isolation of secreting cells with signal proportional to their respective FACS gating (Figure 5B). The selected sub-populations were sorted into a 96-well plate and expanded out of the particles over the course of ∼10 days (Figure 5C). Samples were then plated at the same cell density and bulk antibody production of the different subpopulations was measured by ELISA and compared with non-sorted control samples. We observed a 26% increase in total IgG production for the sorted sub-population 1 (all secreting cells) in comparison to the pre-sort control population (Figure 5D). This increase is most likely due to the removal of cells that are no longer secreting IgG following the sort, which was measured to be ∼25% based on microscopy analysis (Figure 4C) as well as flow analysis (Figure 5A). For sorted sub-population 2 (the top 20% of secretors) we measured an average increase of 41% in total IgG production (n =4) with a maximum increase among the samples of 58% (Figure 5D), indicating the capability of the platform to select out functionally higher producing sub-populations that maintain the phenotype for at least 10 days.

**Figure 5.**
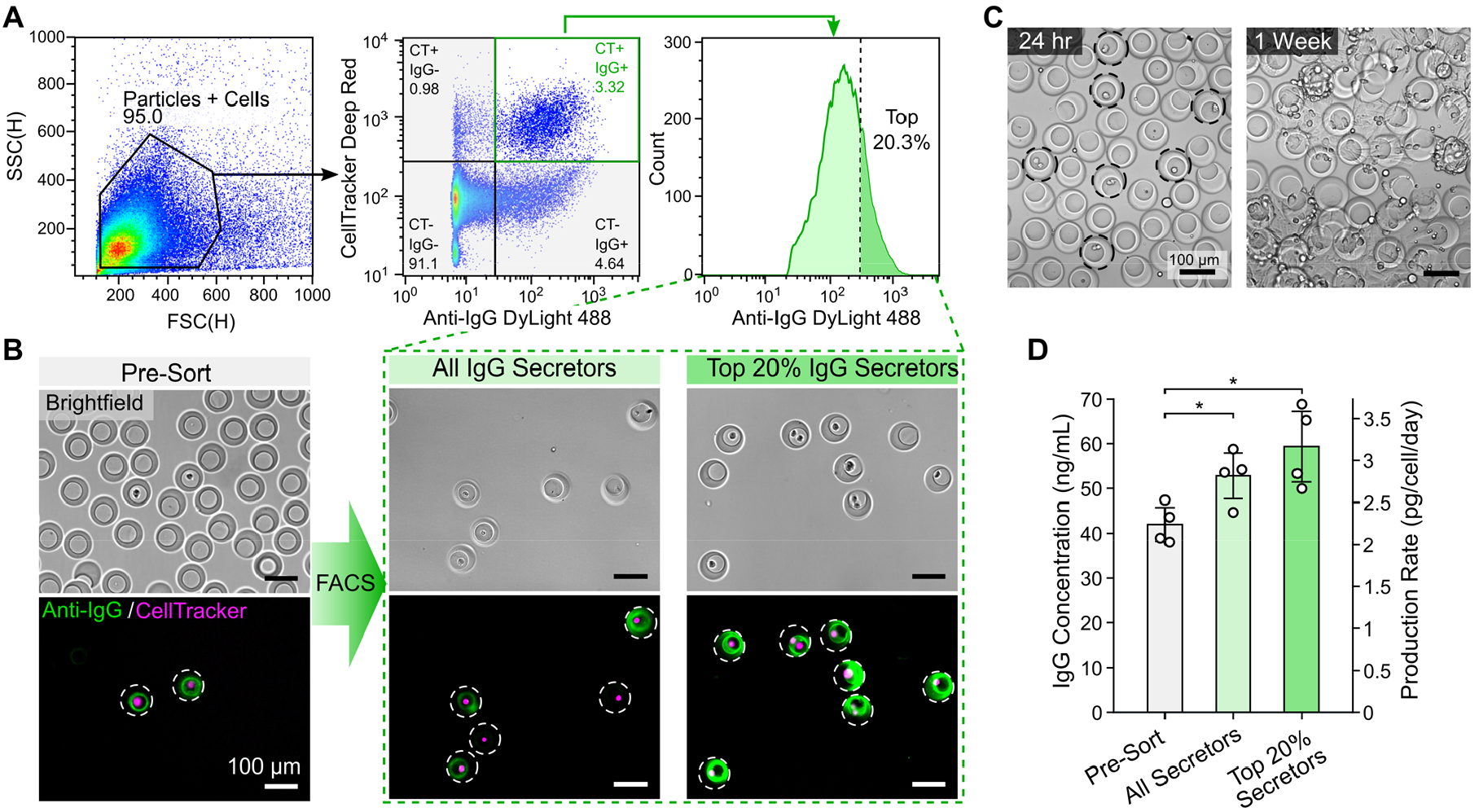
Selection of highly secreting cell sub-populations using FACS. **(A)** The secreting population of cells were gated and sorted based on IgG secretion signal and CellTracker™ (CT) labelling. Both cells with any secretion signal disparate from background as well as the top 20% of secretors were separately sorted. **(B)** Microscopy images before and after sorting show enrichment of secreting cells with fluorescence intensity proportional to the selection criteria. **(C)** Example images of cells expanded out of nanovials after encapsulation and release. **(D)** After expanding isolated cells for 10 days, bulk ELISA was performed to determine the production rate for the different sub-populations. The full experiment was performed with separate cell passages (n = 4). Statistical significance based on standard two-tailed *t*-test (*p<0.05). Error bars represent s.d.

### Isolation of antigen-specific IgG secreting cells

In a final application, we tested the feasibility of using our nanovial platform to screen and isolate antigen-specific antibody producing cells out of a background of similar, non-specific, antibody producers (Figure 6, Figure S9). Rather than selecting simply for high secretion rates, this type of assay replicates strategies taken for antibody discovery against a novel target antigen using our nanovial secretion screen. HyHel-5 hybridoma cells secreting anti-hen egg lysozyme (anti-HEL) antibodies (IgG1) were used as our target cell population and were diluted into a background of similar, CellTracker™ Blue (CTB) labeled, 9E10 hybridoma cells producing anti-myc antibodies (IgG1).

**Figure 6.**
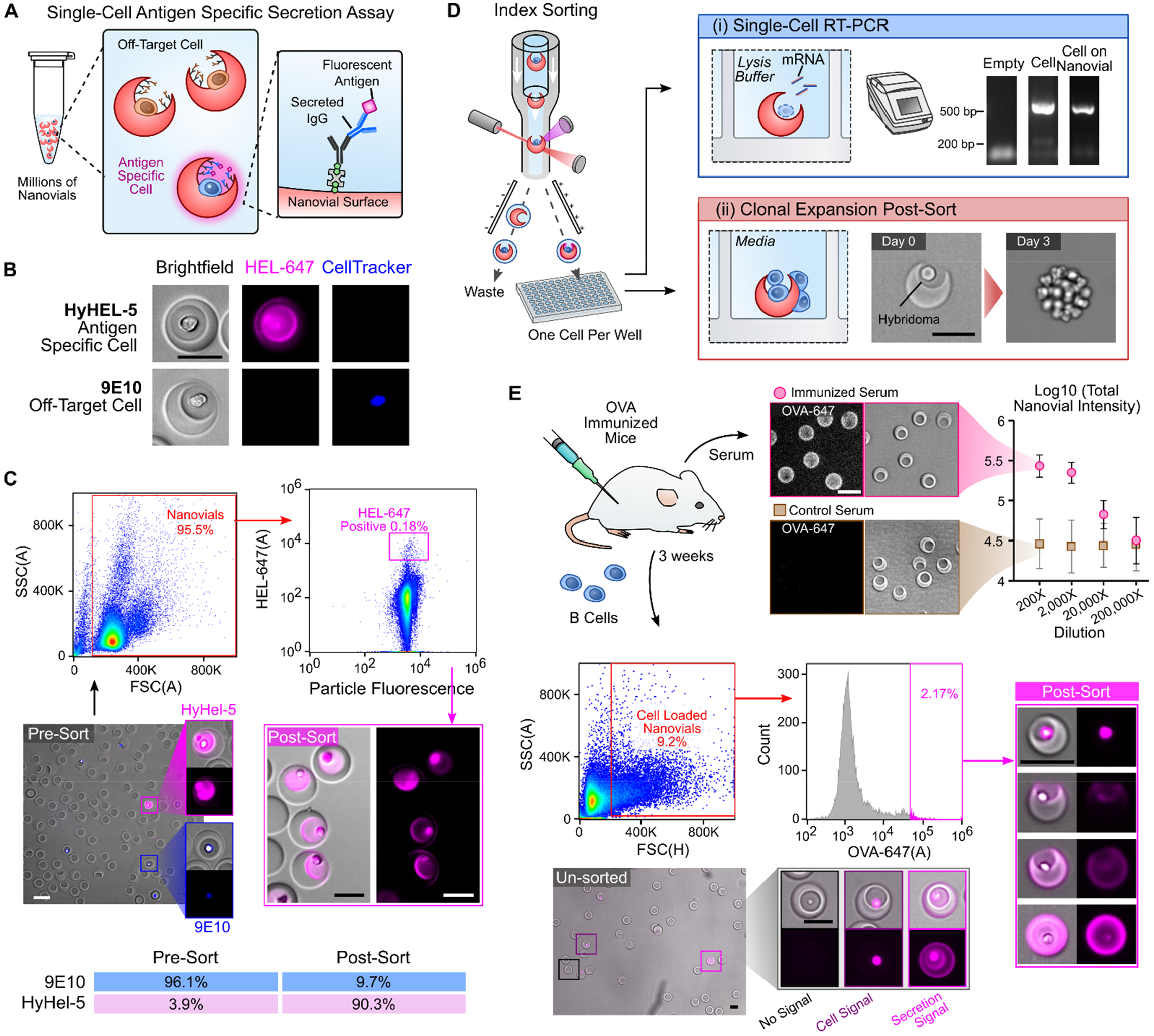
Detection and sorting of antigen-specific antibody secreting cells. **(A)** Hybridomas or B cells loaded into nanovials secrete antibodies which are captured onto the nanovial surface via biotinylated capture antibodies. Captured IgG is then labeled with fluorescent antigen to assess specificity of the secreted IgG. **(B)** Fluorescence microscopy images reveal specific signal from binding to Alexa Fluor™ 647 conjugated hen egg lysozyme (HEL-647) on nanovials containing a hybridoma line (HyHEL-5) secreting IgG HEL while a cell line (9E10) secreting an off-target IgG results in no signal. **(C)** HyHel-5 hybridomas spiked into 9E10 hybridomas (CellTracker™ Blue) at a 1:25 ratio were assayed by fluorescence microscopy for HEL-specific signal before FACS sorting (Pre-Sort) and after sorting using a high HEL-AF647 gate (Post-Sort). A table showing the Pre-Sort and Post-Sort statistics is shown. **(D)** Using the index sorting function of the flow cytometer we sorted single hybridomas loaded onto nanovials and (i) demonstrate that RNA can be reverse transcribed and amplified from single cells loaded on nanovials at comparable rates to single cells sorted that were freely suspended. A gel band corresponding to the correct heavy chain amplicon length is observed. (ii) We additionally demonstrate that individual hybridomas can be expanded into a clonal colony directly from the nanovials following sorting. **(E)** Antibody-secreting B cells from ovalbumin (OVA)-immunized mice are assayed for antigen specificity using nanovials. OVA-specific signal is observed directly on the nanovials by fluorescence microscopy down to 20,000-times dilution of serum from OVA-immunized mice but not control mice. B cells loaded on nanovials were observed to have both cell surface signal and OVA-specific signal by fluorescence microscopy. Following FACS sorting based on a gate encompassing the top 2.17% of OVA fluorescence area signal, OVA-specific antibody-secreting cells were successfully imaged possessing minimal cell surface staining with OVA, along with some cells staining strongly but possessing no OVA-specific secreted antibody signal.

Mixed populations of HyHel-5 and 9E10 hybridoma cells were loaded and bound to nanovials using anti-CD45 cell capture antibodies as described previously, and all antibodies secreted from both cell types were captured non-specifically onto the nanovial surfaces using biotinylated secretion capture antibodies targeting mouse IgG heavy and light chains (Figure 6A). Following incubation to capture secreted IgG, we labeled particles with fluorescently conjugated HEL antigen (HEL-647), revealing the presence of only the antigen-specific antibody secretions (**Figure 6A**). When assaying mixed hybridoma populations, this assay format yielded strong and specific signal with over 80% of nanovials containing HyHel-5 hybridoma cells staining strongly for antigen specific secretion after one hour, while no detectable signal was present on nanovials containing 9E10 cells (**Figure 6B**). Over one million nanovial events were analyzed and a gate encompassing 0.18% of the nanovial population was sorted based off of their high HEL fluorescence area. Subsequent microscopic analysis confirmed highly enriched populations of strongly labeled HEL-647^+^/CTB^-^ cell-loaded nanovials in the post-sort population (>90% purity), confirming that the entire analysis and sorting workflow was compatible with detecting antigen-specific antibody produced by target suspension cells (Figure 6C). In addition to bulk recovery of enriched cell samples producing antigen-specific antibodies, we recovered single hybridoma-loaded nanovials directly into individual wells of a 96-well plate for downstream sequence recovery and re-growth (Figure 6D). We were able to lyse individual cells loaded on nanovials that were deposited into wells and amplify antibody sequence information through standard single-cell RT-PCR (Figure 6D, Figure S10) with similar efficiency (∼50%) compared to freely suspended single cells, indicating that nanovial materials do not interfere with the RT-PCR chemistry. We were also able to sort single hybridoma clones loaded on nanovials into individual wells, culture them and expand them into clonal sub-populations (Figure 6D).

The antigen binding assay was extended to a second antigen and to primary B cells from immunized mice. We first confirmed that ovalbumin (OVA)-immunized mice produced IgG that was OVA-specific by incubating nanovials in serum from either immunized or control mice. Anti-IgG coated particles incubated in serum of immunized mice and exposed to fluorescent OVA yielded differentiable fluorescence signal down to a 20,000-times serum dilution, whereas no detectable signal was generated from control serum at any concentration (Figure 6E, Figure S11). We then loaded and sorted OVA-specific antibody-secreting B cells isolated from the spleens of immunized mice (Figure 6E). A subset of the analyzed and sorted cell-containing nanovials was associated with strong OVA-specific signal on the nanovials themselves, indicating capture of secreted antibodies (Figure 6E). There was little to no surface stain on the captured cells (Figure 6E), suggesting we isolated plasma B cells which canonically lack high levels of B cell receptor (BCR) on the cell surface. Another subset of the B cells acquired strong surface fluorescence after exposure to fluorescent antigen, but did not secrete detectable levels onto the nanovial surface, suggesting direct binding to BCRs, or non-specific binding to membrane compromised cells. A third population of cells had both cell surface BCR staining and staining on the nanovial surfaces, indicative of secreted antibodies. The top ∼2% of nanovials with the highest OVA staining were sorted by FACS and visualized, yielding the same subsets of cell types (secretion signal, cell surface signal, and combined secretion and cell surface signal) that likely reflects the ability to isolate a larger repertoire of B cell populations, including plasma B cells difficult to isolate with standard antigen baiting workflows.

## Discussion

The nanovial platform provides specific advantages in sorting cells based on secretions, and also lays the foundation for the next generation of single-cell and single molecule assays using existing accessible instrumentation. Two key features of the nanovial platform allow for widely accessible analysis and sorting of cells based on secretions: (i) the ability to form uniform compartments containing single cells in small volumes with minimal crosstalk using simple pipetting and no devices; and (ii) the compatibility of the nanovials with commercially available flow sorters such that viable cells can be sorted based on their associated secretions. Nanovials allow for seeding of cells into their open cavities and emulsification to form hundreds of thousands to millions of sealed compartments all in parallel using simple pipetting operations. Parallel encapsulation in < 1 min is an important differentiator from microfluidic techniques, which often require tens of minutes to several hours to sequentially form droplets while cells remain mixed and secrete within the input sample volume (*37*). Rapid emulsification minimizes cross-talk by minimizing the time cells secrete into the input sample volume (e.g. a syringe) before they are encapsulated which is critical for enabling screening of large cell populations. Parallel encapsulation also ensures that the secretion assay starts at the same time point, reducing differences in secretion signal resulting from different secretion times that can add noise, reducing selection accuracy. Nanovial emulsification and analysis steps can also be more easily performed in a sterile environment (e.g. biosafety hood) using standard sterile plasticware (e.g. well plates, pipette tips), as opposed to bulkier equipment. Spherical gel particles have been used previously to template emulsions (*28, 29*), however, the lack of a cavity precluded their use with mammalian cells, and no secretion assays or compatibility with flow cytometry were demonstrated. We have also previously used amphiphilic cavity-containing particles to form uniform emulsions and perform molecular and cellular assays (*25*–*27*); however, these previous systems were not compatible with commercial flow cytometers due to their hydrophobic components.

Although microfluidics is used to manufacture the nanovials, the particles can be easily produced in batch (>10 million nanovials per batch, (Figure S1) and provided to other labs that do not have expertise in microfluidics. Shipping of particles is far more cost-effective and rapid than replicating microfluidic setups for generation and sorting of droplets. Further, the amount of additional expertise required to use the nanovial system is substantially lower than required for microfluidic-based approaches due to the familiar handling steps. Collaborators across the globe have been able to perform nanovial assays without any hands-on training, including the experiments shown in Figure S8 which were performed solely at Johns Hopkins University in a protein engineering lab without any access to microfluidic devices.

The nanovial system is also compatible with commercial flow cytometers which enabled us to screen over a million nanovials in under an hour across the experiments described in this report (at rates exceeding 600 events/second). This throughput outperforms previous droplet microfluidic sorters applied to antibody discovery workflows (*15*). Throughput can be further improved by reduction of the nanovial size for compatibility with smaller FACS nozzle sizes compatible with sorting rates exceeding 10,000 events per second. Further, incorporation of more scalable enrichment techniques such as magnetic or density driven separation could enable throughputs exceeding 10 million cells per day. Besides increasing availability to labs without expertise in microfluidics or specialized commercial instruments, there are also unique advantages of this new approach for encapsulation of mammalian cells.

The structure and surface of the nanovials provide unique capabilities for tuning the number and type of captured cells, analysis of adherent cell secretions, and analysis and sorting of clonal colonies. The size and opening diameter of the cavity within a nanovial can be tuned (Figure S2), enabling more deterministic loading of single cells based on size exclusion effects (*38*). This could allow more rapid screening because cell loading rates are not dictated by Poisson statistics. Besides structural changes to the particles, the surface of particles can also be functionalized with affinity agents that specifically enrich certain populations of cells, such as antibodies to CD3 that enrich T cells from a mixed population. The surface of the particle also enables the attachment and growth of adherent cell populations, such as the CHO and HEK293 cells used herein. Analysis of secretions from adherent cells can be challenging with other microfluidic techniques which require cells to be in suspension to flow into devices and sort afterwards. Adherent cells begin to undergo apoptosis when remaining in suspension and it is expected that secretion rates of biomolecules would change in this condition (*39*). The inability to perform these assays has led to a dearth of information on the secretion phenotypes and heterogeneity of adherent cells with important secretion products in health and disease, such as mesenchymal stem cells, glandular epithelial cells, endothelial cells, glial cells, tumor cells and even adherent bacterial biofilms. Further, in the nanovial system, when single cells are initially seeded onto particles they adhere and grow without nutrient limitations (Figure S6). The clonal colonies can then be encapsulated to form dropicles, enabling high-throughput screening based on the combination of growth and per cell secretion (i.e. overall biomass produced per unit time), potentially overcoming previous challenges in cell line selection in which growth and biologic production can be in a trade-off relation (*13*).

Sorting based on secretions extends beyond selection of high antibody-titer cell lines to many applications of importance in life sciences and biotechnology. Discovering high affinity antibody therapeutics relies on the selection of B and plasma cells producing antibodies with high affinity amongst a large background of clones (*7*). Although direct antigen binding to cell surface-expressed BCRs followed by FACS has been used to isolate antigen-specific B cells, plasma cells lose BCR expression, but represent an important population of antibody-secreting cells that have gone through affinity maturation, and may yield higher affinity antibody sequences (*40*). In addition, highly-secreting plasma cells and plasmablasts have larger numbers of transcripts encoding the antibody sequence, making them prime targets for sequence recovery by single-cell RT-PCR or downstream single-cell sequencing. More generally, the activity of immune cells is largely connected to their secretion profiles, which direct communication and effector functions and can be better studied by sorting out specific sub-populations for further functional testing in vivo or in vitro. The effectiveness of chimeric antigen receptor-T cell batches also appears to depend on a multifunctional secretion of cytokines, such that sorting populations based on secretion profiles may enhance therapeutic activity (*41, 42*). Finally, processes of directed evolution of cell products and cells themselves can benefit from larger numbers of clones being screened, mutagenized, and expanded across multiple cycles using an efficient process relying on standard equipment and expertise (*43*).

In sum, our new approach to encapsulate single entities into uniform compartments with a solid phase can democratize access to cutting-edge assays, creating a modular “lab on a particle” platform for a number of single-cell and single-molecule assays. Encapsulation is a key component for single-cell nucleic acid sequencing. Clonal colonies of bacteria, yeast, or algae producing engineered proteins (e.g. fluorescent proteins) can also be maintained in nanovials and sorted based on desirable features (e.g. intensity at particular excitation/emission wavelengths). Moving beyond cells, the compartments formed can enable digital nucleic acid amplification assays and immunoassays, where the solid phase provides potential for barcoding and capturing of amplified assay signals. Given the ability to rapidly deploy our approach with established lab infrastructure we anticipate widespread applications of lab on a particle technology across a range of these single-cell and single-molecule assays in the near future. Rapid deployment across the world is of particular importance during emerging pandemics, enabling collective distributed research and development, and removing the bottlenecks created by the sparsity of skilled groups able to contribute to key points in therapeutic and diagnostic pipelines.

## Methods

### List of reagents and resources

| REAGENT OR RESOURCE | SOURCE | IDENTIFIER |
| --- | --- | --- |
| **Antibodies** |  |  |
| Goat anti-human IgG H&L, Dylight® 488 conjugated | Abcam | ab96911 |
| Goat anti-human IgG Fc, biotin conjugated | Thermo Fisher | A18821 |
| Rat anti-mouse CD45, biotin, Clone: 30-F11 | Thermo Fisher | 50-115-49 |
| Biotin anti-mouse CD19, Clone: 6D5 | Biolegend | 115503 |
| Goat anti-mouse IgG H&L, biotin | SouthernBiotech | 1034-08 |
| **Chemicals, Peptides and Recombinant Proteins** |  |  |
| *Nanovial fabrication and modification* |  |  |
| 4-arm Poly(ethylene glycol) norbornene terminated | Sigma-Aldrich | 808474-1G |
| Dithiothreitol | Sigma-Aldrich | 10197777001 |
| Lithium phenyl-2,4,6-trimethylbenzoylphosphinate | Sigma-Aldrich | 900889-1G |
| Biotin PEG thiol | Nanocs | PG2-BNTH-5k |
| RGD peptide (Ac-RGDSPGERCG-NH<sub>2</sub>) | Genscript | Custom Order |
| Dextran from Leuconostoc spp. (40,000) | Sigma-Aldrich | 31389-25G |
| Hexane, mixture of isomers | Sigma-Aldrich | 178918-2.5L |
| 1H,1H,2H,2H-Perfluoro-1-octanol | Sigma-Aldrich | 370533-5G |
| Thiol-Poly-L-Lysine-Thiol | Nanosoft Polymers | 12570-32,000 |
| Fibronectin Human Plasma, Protein | Thermo Fisher | 33016015 |
| Pico-Surf(5%(w/w)) in Novec™ 7500 | Sphere Fluidics | CO24 |
| 3M Novec™ 7500 | Gallade Chemical | N17501 |
| Mineral oil, light | Sigma-Aldrich | 330779-1L |
| *Cell Culture* |  |  |
| Fetal Bovine Serum US Certified | Invitrogen | 16000044 |
| Penicillin-Streptomycin (10,000 U/mL) | Invitrogen | 15140122 |
| DMEM, high glucose, pyruvate | Invitrogen | 11995065 |
| RPMI | Invitrogen | 11875093 |
| IMDM | Invitrogen | 12440053 |
| F-12 | Invitrogen | 11765054 |
| L Glutamine | Invitrogen | 25030081 |
| Insulin, Human recombinant | Sigma-Aldrich | 91077C-100MG |
| Trace Elements A | Thermo Fisher | MT99182CI |
| Trace Elements B | Thermo Fisher | MT99175CI |
| Methotrexate hydrate | Sigma-Aldrich | M8407-100MG |
| Gibco™ ExpiCHO™ Expression Medium | Fisher Scientific | A2910001 |
| *Secretion Assay* |  |  |
| Streptavidin (5 mg/mL) | Thermo Fisher | 434302 |
| Streptavidin, Alexa Fluor™ conjugated | Fisher Scientific | S11226A |
| CellTracker™ Deep Red Dye | Thermo Fisher | C34565 |
| CellTracker™ Blue CMAC Dye | Thermo Fisher | C2110 |
| DPBS (Ca+,Mg+) | Invitrogen | 14040133 |
| BSA Bovine Serum Albumin (BSA), Fraction V—Low-Endotoxin Grade | GeminiBio | 700-102P |
| Pluronic™ F-127 | Sigma | P2443-250G |
| CellTrics™ Filters, 20 µm | Thermo Fisher | NC9699018 |
| Reversible Strainers (37 µm) | Stemcell Technologies | 27215 |
| Lysozyme (Hen Egg White) | Aviva Systems Biology | OORA00201 |
| Alexa Fluor™ 647 Microscale Protein Labeling Kit | Thermo Fisher | A30009 |
| Ovalbumin, Alexa Fluor™ 647 | Thermo Fisher | O34784 |
| *Immunization* |  |  |
| Imject™ Alum Adjuvant | Thermo Fisher | 77161 |
| Ovalbumin | Biosearch Technologies | N-5051-10 |
| EasySep™ Mouse B Cell Isolation Kit | STEMCELL™ Technologies | #19854 |
| **Commercial Assays** |  |  |
| IgG (Total) Human ELISA Kit | Invitrogen, | BMS2091 |
| **Experimental Models: Cell Lines** |  |  |
| Hamster: CHO-K1 (CCL61) | ATCC | CCL61 |
| Hamster: CHO DP-12 (CRL-12445) | ATCC | CRL-12445 |
| Human: HEK293 cells | ATCC | CRL-1573™ |
| Hamster: ExpiCHO-S cells | Thermo Fisher | A29127 |
| Mouse: HyHEL-5 | Richard Willson |  |
| Mouse: 9E10 | ATCC | CRL-1729™ |
| **Experimental Models: Organisms/Strains** |  |  |
| Mouse: C56BL/6J mice | Jackson Laboratory |  |
| **Single-Cell RT-PCR** |  |  |
| RNasin® Plus Ribonuclease Inhibitor | Promega | N261B |
| Phusion High Fidelity DNA Polymerase | Thermo Fisher | F-549S |
| **Software and Algorithms** |  |  |
| Image J | NIH | https://imagej.nih.gov/ij |
| Matlab | Mathworks | Version 2016b |
| FlowJo | FlowJo, LLC |  |

### Fabrication of nanovials

Nanovials were fabricated using a standard PDMS microfluidic flow focusing droplet generator. A PEG phase comprised of 28.9% w/w 4-arm PEG-norbornene (Sigma), 3% w/w LAP (lithium phenyl-2,4,6-trimethylbenzoylphosphinate, Sigma), and 1 mg/mL biotin-PEG-thiol (5000 MW, Nanocs) in phosphate buffered saline (PBS, pH 7.2) was co-injected with a dextran phase comprised of 11% w/w 40 kDa dextran (Sigma), 1.3% w/w DTT (dithiothreitol, Sigma), and 5 mM RGD peptide (Ac-RGDSPGERCG-NH2, Genscript) in PBS at a rates of 0.5 - 5 µL/min, depending on nanovial size, using syringe pumps (Harvard Apparatus PHD 2000). An oil phase comprised of Novec™ 7500 (3M) and 0.25% w/w fluorinated surfactant was injected at a rate of 10 - 42 µL/min to partition the aqueous phases into monodisperse water in oil droplets. PEG and dextran polymers phase separated on chip after approximately 5 seconds. The PEG phase was crosslinked with focused UV light through a DAPI filter set and microscope objective (Nikon, Eclipse Ti-S) near the outlet region of the microfluidic device. Crosslinked nanovials were collected and oil and dextran were removed using a series of washing steps. Briefly, excess oil was removed by pipetting and a layer of PBS was added on top of the remaining emulsions. A solution of 20% v/v perfluorooctanol (PFO, Sigma) in Novec™ 7500 was then added to destabilize the emulsions and transfer nanovials to the PBS phase. Excess oil was removed and samples were washed two times with Novec™ 7500 to remove remaining surfactant. Novec™ 7500 was removed by pipetting and residual oil was removed by washing two to three times with hexane (Sigma). Samples were then washed three times with PBS to remove dextran from the system. For cell experiments, nanovials were sterilized by incubating in 70% ethanol overnight. Nanovials were then washed five times with washing buffer comprised of 0.05% Pluronic™ F-127 (Sigma), 1% penicillin/streptomycin (Invitrogen), and 0.5% bovine serum albumum (BSA, GeminiBio) in PBS and stored in a conical tube at 4°C.

#### PLL modification of nanovials

To modify the nanovial surface with poly-L-lysine (PLL), a 100 µL volume of concentrated 55 µm nanovials was suspended in 1 mL solution consisting of 1 mg/mL thiol-poly-L-lysine-thiol (32 kDa, Nanosoft Polymers) and 0.2 % w/w LAP. The nanovial suspension was transferred to a glass vial pre-coated with Sigmacote (Sigma SL2-25ML) and the vial was placed on top of a mini stir plate (IKA Lab Disc IKAMAG Magnetic Stirrer) with a micro stir bar in it. With continuous mixing, the particle suspension was exposed to UV for 60 seconds at a power of 2.9 mW/cm^2^. The particle suspension was retrieved into a 1.5 mL Eppendorf tube and washed with PBS containing 0.05% Pluronic™ F-127 three times. To further modify PLL-conjugated nanovials with fibronectin, 25 µL of concentrated nanovials were incubated in 500 µL of 10 µg/mL or 500 µg/mL fibronectin solution for one hour at room temperature and washed with washing buffer three times.

### Nanovial handling and modification

#### Washing buffer to coat surfaces and wash nanovials

To reduce loss of nanovials due to sticking, all pipette tips, serological pipettes, microcentrifuge tubes, conical tubes, FACS tubes, and well-plates were pre-coated with sterile washing buffer comprised of 0.05% Pluronic™ F-127 (Sigma), 1% penicillin/streptomycin (Invitrogen), and 0.5% bovine serum albumin (BSA, GeminiBio) in PBS unless otherwise noted.

#### Nanovial assay washing procedure

In general, nanovials are washed by centrifuging at 300 g for 2 – 5 min to create a pellet. Supernatant is removed by carefully pipetting or aspirating fluid above the pellet. Nanovials are then re-suspended in washing buffer or media at a 10-fold dilution. This procedure is repeated as needed.

#### Streptavidin labeling

Biotinylated nanovials are pelleted and supernatant is removed. For 35 and 55 µm nanovials, samples were reconstituted at a five times dilution in washing buffer containing 60 µg/mL of streptavidin and incubated for 15 – 30 min at room temperature. For fluorescent staining of nanovials, 0.6 – 6 µg/mL fluorophore-conjugated streptavidin was included in addition to the unlabeled streptavidin. For 85 µm nanovials, 400 µg/mL streptavidin solution was added to the concentrated pellet at a 1:1 ratio and pipetted to mix evenly. Fluorophore-conjugated streptavidin was prepared at 4 – 40 µg/mL. Excess streptavidin was removed by washing three times.

#### Biotinylated antibody labeling

Streptavidin coated nanovials are pelleted and supernatant is removed. For 35 and 55 µm nanovials, samples were reconstituted at a five times dilution in washing buffer containing 5 - 20 µg/mL of the desired biotinylated antibody or mixture of antibodies and incubated for 15 – 30 min at room temperature to allow for binding to streptavidin. For 85 µm nanovials, a working solution containing 30 µg/mL of biotinylated antibodies was added to the concentrated pellet at a 1:1 ratio and pipetted to mix evenly.

### Cell culture

All cells were cultured in incubators at 37°C and 5% CO_2_ in static conditions unless otherwise noted.

#### Chinese hamster ovary (CHO Cells)

CHO DP-12 cells (ATCC, CRL-12445™) were maintained according to manufacturer’s specifications. Cell culture media was comprised of DMEM (Invitrogen) supplemented with 10% fetal bovine serum (FBS, Invitrogen), 1% penicillin/streptomycin, 0.002 mg/mL recombinant human insulin (Sigma), 0.1% Trace Elements A (Fisher Scientific), 0.1% Trace Elements B (Fisher Scientific), and 200 nM Methotrexate (MTX, SIGMA). CHO-K1 cells (ATCC, CCL61™) were cultured in F-12 base media (Invitrogen) supplemented with 10% FBS and 1% penicillin/streptomycin (Invitrogen). ExpiCHO cells (Fisher Scientific A29127™) were cultured in ExpiCHO™ Expression Medium (Fisher Scientific) on an orbital shaker in a 37°C incubator with ≥80% relative humidity and 8% CO_2_. The shake speed was set to 120 rpm with 25 mm shaking diameter. Cells were seeded at 0.2 × 10^6^ viable cells/mL and subcultured when the cell density reached 4 × 10^6^– 6 × 10^6^ viable cells/mL.

#### Human embryonic kidney (HEK) cells

HEK 293T cells (Thermo Life Technologies) were cultured in DMEM medium with 10% FBS, L-glutamine (2 mM), and penicillin-streptomycin (500 µg/mL).

#### Hybridoma cells

HyHel-5 cells were maintained in IMDM media (Invitrogen) supplemented with 10% FBS (Invitrogen) and 1% penicillin/streptomycin (Invitrogen). 9E10 cells (ATCC, CRL-1729™) were maintained in RPMI media (Invitrogen) supplemented with 10% FBS (Invitrogen) and 1% penicillin/streptomycin (Invitrogen). Cells were passaged down to a final concentration of 4×10^5^ cells/mL every two days.

### Mouse immunization, serum isolation, and splenocyte isolation

All experiments involving animals, animal cells, or tissues were performed in accordance with the Chancellor’s Animal Research Committee ethical guidelines at the University of California Los Angeles under protocol no ARC-2015-125. Ten week old C56BL/6J mice (Jackson Laboratory) were immunized for a total of 7 times over 28 days with recombinant ovalbumin (Biosearch Technologies) as a model antigen. Immunizations were made by mixing ovalbumin at 1 µg/µL in PBS with equal volume of Imject™ alum adjuvant (ThermoFisher Scientific). 250 µg and 25 µg doses of ovalbumin were used for initial immunization, and subsequent 7 boosters, respectively. Four days after final immunization, mice were euthanized with isoflurane overdose followed by cervical dislocation and sterilized by spraying with 70% ethanol. for tissue and blood collection. For splenocyte isolation, the spleen was removed, cut into small pieces with scissors, and pushed through a 70 µm cell strainer in a 10 cm petri dish containing 10 mL cold PBS, using the plunger of a 10 mL syringe. The same syringe was then used to dissociate tissue clumps by drawing and flushing the solution through a 25-gauge needle 3 times. Cells were pelleted by centrifugation at 300 g for 10 min and resuspended at 10^8^ cells/mL in EasySep™ buffer and B lineage cells were isolated using an EasySep™ mouse B cell negative isolation kit (StemCell Technologies, #19854). For serum, blood was collected directly from left ventricle via cardiac puncture immediately after euthanasia. Blood was allowed to clot at room temperature for 4 hours, before centrifuging at 10,000 g for 10 min to separate serum.

### Cell loading in nanovials

#### Nanovial well-plate seeding

Wells were first partially filled with media or buffer. Diluted nanovials were then transferred into the well using a micropipette or serological pipette and dispersed by pipetting up and down in circular motions. Nanovials were then allowed to settle to the bottom of the well for 15 – 60 min, with longer times being required for smaller nanovials. A diluted cell suspension was then carefully pipetted into each well being careful to not disturb the nanovials. Cells were then allowed to settle for 15 – 30 min and then transferred to an incubator unless otherwise noted. The amount of nanovials and cells to add into each well was determined both analytically and experimentally. General values for different nanovial sizes are shown in Table 1.

**Table 1.** Volume of concentrated nanovials to add for different well and nanovial sizes.

| Well Plate Size | 35 $\mu\text{m}$ Nanovials | 55 $\mu\text{m}$ Nanovials | 85 $\mu\text{m}$ Nanovials |
| --- | --- | --- | --- |
| 24-well | 6 $\mu\text{L}$ | 8 $\mu\text{L}$ | 15 $\mu\text{L}$ |
| 12-well | 12 $\mu\text{L}$ | 16 $\mu\text{L}$ | 30 $\mu\text{L}$ |
| 6-well | 29 $\mu\text{L}$ | 40 $\mu\text{L}$ | 67 $\mu\text{L}$ |

#### Characterization of cell loading in nanovials

Nanovials were prepared using the modification procedures described above. 85 µm, 55 µm, and 35 µm nanovials were used for CHO DP-12, ExpiCHO/HEK293, and B cells, respectively. RGD, RGD+PLL, and RGD+PLL+ Fibronectin modified nanovials were prepared following procedures described above. Antibody-coated nanovials were prepared follow procedures above using 8 µg/mL of aCD45-biotin, 8 µg/mL of aCD19-biotin, or 8 µg/mL aCD45-biotin + 8 µg/mL of aCD19-biotin. For all conditions, modified nanovials were seeded into media-containing wells following procedures described above. Cells were fluorescently labeled using CellTracker™ following the manufacturer’s specification and seeded into the nanovials by pipetting. HEK293 cells were first biotinylated using EZ-Link Sulfo-NHS-SS-Biotin (ThermoFisher) following the manufacturer’s specified procedure. B cells were obtained from mice following procedures described in ‘Mouse immunization, serum isolation, and splenocyte isolation’. CHO DP-12 and ExpiCHO cells were incubated for 2-4 hrs in an incubator at 37°C to allow for cell binding. HEK293 cells and B cells were incubated for 1 hr at 37°C to facilitate binding. After incubating, samples were transferred by pipetting and unbound cells were removed from the nanovial solution by running through a cell strainer. For 85 µm nanovials a 37 µm cell strainer was used (Stemcell Technologies) and for 35 and 55 µm nanovials a 20 µm cell strainer was used (CellTrics, ThermoFisher). Nanovials and cell-containing cells were recovered by inverting the cell strainers over a conical tube and passing washing buffer through it with a pipette. Samples were then imaged in well plates using bright field and fluorescence microscopy. Cell loading fraction was characterized using either a Matlab algorithm (CHO DP-12, ExpiCHO, B cells), or using flow cytometry (HEK293, CytoFlex, Beckman Coulter).

### Dropicle formation and characterization

Nanovials with 85 µm diameters were suspended in DMEM base media (Invitrogen) and then pelleted and supernatant removed by pipetting. An oil phase comprised of Novec™ 7500 and 2% w/w PicoSurf was added to the particle suspension at approximately two times the remaining volume. The sample was then vigorously pipetted for 30 - 60 s (∼100 pipettes) using a 200 µL micropipette (Eppendorf). The resulting water-in-oil emulsion was then carefully transferred into a PDMS reservoir by pipetting and imaged. Size distribution characterization was then performed using image analysis algorithms in MATLAB. For the fluorescent images shown in Figure 1B, particles were first labeled with AlexaFluor™ 568 streptavidin and suspended in PBS containing 2 mg/mL fluorescein isothiocyanate-dextran (500,000 MW, Sigma). For 35 and 55 µm nanovials the same procedure was used except the nanovials were first diluted 10-fold to reduce fraction of aggregates formed.

### Dropicle emulsion breaking

To recover cells back into an aqueous phase following dropicle formation, excess oil was first removed via pipetting and several mL of media was added on top of the emulsions. To destabilize the emulsions a destabilizing agent (20% v/v PFO in Novec™ 7500) was pipetted on top of the emulsion layer and the sample was gently agitated. After 5 min most of the droplets were merged and nanovials and any associated cells were transferred into the bulk media phase. Optionally, samples can be centrifuged for 15-30 s at 200 g to coalesce remaining droplets.

### Energy minimization theory

The volume energy curve depicted in Figure S5 was calculated using our previously reported approach (*35*), which assumes that the energy of the system is dominated by surface energy. In Figure S5A, for each given fluid volume, the fluid assumes a morphology to minimize the energy of the system. In Figure S5B, for the two-particle system we calculate the fluid distribution that minimizes the total energy of the system of two particles containing two separate fluid volumes, with the modified assumption that below a normalized volume V/V_0_ =2, the particles remain as aggregates. We assume that the interfacial tension between PEG and water is negligible such that the normalized interfacial tensions are

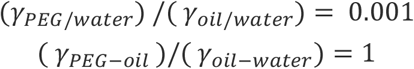

where *y*_*PEG/water*_ is the interfacial tension between the PEG based nanovials and water, *y*_*oil/water*_ is the tension between oil and water and *y*_*PEG/oil*_ is the tension between the nanovials and oil.

### Cell viability characterization

For viability studies depicted in Figure S6, CHO DP-12 cells were seeded into 85 µm RGD modified nanovials. As a control, free cells were seeded into a separate well plate containing no nanovials. Cells were allowed to adhere for 4 hours. Samples containing both the cells and nanovials were transferred into a 15 mL conical tube, exchanged with fresh media and then concentrated via centrifugation and aspiration. Novec™ + 2% w/w PicoSurf was added to the concentrated sample and pipetted for 30 s to encapsulate the nanovials and associated cells into dropicles as discussed above. Light mineral oil was added on top of the samples to mitigate evaporation during prolonged incubation. Samples were then incubated for 2, 12, and 24 hours in an incubator at 37°C and 5% CO_2_. Dropicle emulsions were broken as described above and the suspension of nanovials and cells were then transferred by pipetting into a separate conical tube. Samples were washed with PBS and then sequentially stained with calcein AM and propidium iodide (live/dead). The control well plate samples were stained directly in the well plates with the live/dead stains. Nanovial samples were transferred back into a well plate, imaged, and then analyzed in MATLAB to determine cell viability statistics.

For cell expansion characterization depicted in Figure S6C-D, samples were prepared as above except cells were first stained with CellTracker™ and then incubated for 2 hrs in a dropicle emulsion state. Recovered nanovials and cells were seeded into wells of a 96-well plate and imaged over the course of a week. A control sample was prepared with cells directly seeded into a well plate after trypsinization.

### CHO cell IgG secretion assay

CHO DP-12 cells producing a human anti-IL-8 antibody were used for these studies. To identify these target cells during downstream analysis CHO DP-12 cells were first stained with CellTracker™ Blue CMAC Dye (Thermo Fisher). 85 µm nanovials containing RGD and cells were seeded into a 12-well plate as described above and then incubated at 37°C for 2 hours to allow cells to adhere to the particles. Nanovials were recovered by pipetting and background cells removed using a 37 µm reversible cell strainer. Recovered nanovials were washed two times with washing buffer and sequentially labeled with streptavidin and biotin goat anti-human IgG Fc (Thermo Fisher, A18821) following standard procedures described above. Samples were then washed and resuspended in CHO DP-12 media. Nanovial samples were then compartmentalized by pipetting with oil and surfactant as described above to create dropicles and incubated for a range of times 0, 1, 2, 4, and 8 hours to allow cells to secrete and to facilitate capture of secreted antibodies onto the associated particle matrix via anti IgG binding sites. After the incubation period, nanovials and associated cells were transferred back into media by breaking the emulsions (see ‘Dropicle emulsion breaking’ section). Samples were then washed, and captured secretions were stained by adding a cell staining buffer (0.05% Pluronic™ F-127, 2% FBS, and 1% penicillin/streptomycin in PBS (Ca^2+^,Mg^2+^)) containing 30 µg/mL Goat anti-human IgG H&L (Dylight® 488, Abcam ab96911) at a 1:1 ratio. After 30 min of staining, samples were then washed three to five times with cell staining buffer and optionally stained with propidium iodide. Samples were imaged in both brightfield and fluorescence channels in a well plate. To characterize secretion amount per particle a MATLAB algorithm was used to identify particles in brightfield and then count the number of cells, check for the presence of dead stain, and integrate the total secretion label fluorescence intensity for each particle.

### Secretion cross-talk analysis experiment

To analyze potential cross-talk when nanovials were not emulsified, samples were prepared as described in the ‘CHO cell IgG secretion assay’ section with several modifications to the protocol. Two sets of samples were prepared: a control sample that was incubated in bulk solution (without emulsification) and a test sample incubated after dropicle formation. Prior to the incubation step a separate suspension of AlexaFluor™ 647 tagged nanovials containing no cells were mixed into the samples. This was done in order to ensure signal on empty nanovials that was measured did not arise from cells that may have detached from the nanovials during various steps of the assay. Samples were washed and modified with biotin goat anti-human IgG Fc as described above. The bulk samples were left to incubate in media while the dropicle samples were emulsified (see ‘Dropicle formation and characterization’). After incubating for 15 hours samples were recovered and washed with washing buffer, stained, and imaged. The amount of cross talk was determined by comparing secretion staining intensity of cell containing nanovials with the intensity on the control particles.

### CHO DP-12 sorting and enrichment from background cells based on secretion

CHO DP-12 cells and CHO-K1 cells were prelabeled with CellTracker™ Deep Red and CellTracker™ Blue (Thermo Fisher). After labelling, cells were mixed together at various ratios (1:5, 1:100, 1:1000) and loaded into nanovials (Seeding density ∼84/mm^2^, lambda ∼0.1). All remaining secretion assay steps were as previously described (see ‘CHO cell IgG secretion assay’). After labelling secretions on samples with goat anti-human IgG H&L Dylight 488, samples were sorted using FACS (BioSorter, Unionbio). Samples were excited using both 488 nm and 561 nm lasers. Events were triggered based on nanovial absorbance from the 561 nm laser. Single nanovial events were gated based on time of flight. Nanovials with secretion signal were sorted by thresholding the peak intensity height collected through a 543/22 nm filter. Samples were sorted directly into a 96-well plate and imaged with a fluorescence microscope to quantify purity and enrichment.

### Analysis of single-cell IgG secretion distributions for transiently transfected cells

HEK293T cells were washed three times with PBS pH 8, and then biotinylated using EZ-Link Sulfo-NHS-SS-Biotin (Thermo Fisher Scientific). Per manufacturer’s protocol, the cells were resuspended at 25×10^6^ cells/mL in PBS pH 8 containing 2 mM of EZ-Link Sulfo-NHS-SS-Biotin reagent and incubated at room temperature for 30 min with rotation. Three washes were then conducted using PBS (pH 7.3) to remove excess byproducts. Cells were then stained with CellTrace™ Violet dye (Thermo Fisher Scientific). Nanovials (55 µm diameter) were prepared by coating with streptavidin and Alexa Fluor™ 647-conjugated streptavidin at a 10:1 ratio following standard procedures (See ‘Nanovial handling and modification’). HEK293T cells were loaded into nanovials as described in ‘Cell loading in nanovials’. After cell loading, nanovials were modified with biotinylated goat anti-human IgG Fc antibody (Thermo Fisher Scientific). Samples were resuspended in DMEM and seeded in a well plate. The anti-programmed death ligand-1 (PD-L1) antibody atezolizumab and the anti-interleukin 8 receptor beta (IL-8Rb) antibody 10H2 were cloned into the gwiz mammalian expression vector (Genlantis) and used for transient transfection of HEK 293T cells loaded into nanovials. Atezolizumab and 10H2 DNA were diluted to 0.05 mg/mL in OptiPro medium (Thermo Life Technologies) with a heavy:light chain ratio of 1:2 and 1:1 respectively, and incubated at room temperature for 5 min. Polyethyleneimine (PEI, Polysciences) was independently diluted to 0.1 mg/mL in OptiPro medium and incubated at room temperature for 5 min. Equal volumes of DNA and polyethyleneimine were mixed and incubated at room temperature for an additional 15 min. The DNA/PEI mixture was then added to the 6-well plate while gently rotating the plate to mix. The plate was allowed to incubate for 16 hrs, emulsified as described in section ‘Dropicle formation and characterization’, and then recovered after another 32 hours (see ‘Dropicle emulsion breaking’). Recovered nanovials were stained with 30 µg/mL goat anti-human IgG H&L DyLight 488 (Abcam) and 1 µg/mL of propidium iodide (Thermo Fisher Scientific) for 30 min at 4°C without rotation. Samples were then washed three times and analyzed via flow cytometry (CytoFlex, Beckman Coulter).

### Enrichment of high IgG producing CHO cells, re-culture, and bulk ELISA on secreted IgG

CHO-DP12 cells were loaded in nanovials and secreted IgG captured as described in the ‘CHO cell IgG secretion assay’ section. After labelling secretions, a fraction of each sample was kept and imaged using fluorescence microscopy. The remaining samples were then sorted using a FACS instrument (On-Chip Sort, On Chip Biotechnologies). Samples were excited with both a 488 nm and 637 nm laser. Particle events were screened based on the forward and side scatter. Particles positive for both cells and secretion signal were gated based on peak fluorescence height collected through a 543/22 nm emission filter and a 676/37 nm emission filter respectively. Two sub-populations were sorted for each sample: (1) particles with cells and positive secretion signal, (2) particles with cells and the top 20% of positive secretion signal. Collected samples were plated and expanded for >10 days. To quantify antibody production of the isolated sub-populations, 30,000 cells from the expanded sub-populations as well as unsorted control samples were plated into a 48-well plate. After cells attached the samples were washed, replaced with 400 µL of fresh media, and incubated for 6 hours. Supernatant was then collected and total human IgG amount was measured using ELISA (IgG (Total) Human ELISA Kit, Invitrogen, BMS2091). Production rate per cell was calculated based off the measured IgG concentration, incubation time, and initial number of cells seeded.

### Hybridoma secretion assay and sorting

Nanovials with 55 µm diameters were used for screening antigen-specific antibody production from HyHel-5 hybridomas. Nanovials were sequentially coated with streptavidin and biotinylated antibodies (20 µg/mL anti-mouse CD45, 20 µg/mL anti-mouse IgG H+L) (see ‘Nanovial handling and modification’). Concurrently 9E10 hybridoma cells were labeled with CellTracker™ Blue dye (Thermo Fisher) and premixed with unlabeled HyHel-5 hybridoma cells at desired concentrations. Importantly the CellTracker™ Blue dye (Thermo Fisher) stain was not used for sorting or enrichment purposes, but served to differentiate the two cell types in pre- and post-sort analysis to benchmark assay performance. The premixed hybridoma cell populations were next loaded into antibody-coated hydrogel nanovials chilled to 4°C (see ‘Cell loading in nanovials’). Nanovials and cells were maintained at 4°C for 1 hour to lower their metabolic activity and prevent accumulation of secretion onto neighboring unloaded nanovials during the binding process. Following adhesion, unbound cells were removed from the sample by filtering through a 20 µm cell-strainer, and the recovered cell-loaded particles were incubated in IMDM media at 37°C for another hour to accumulate secreted products. We found that for rare populations of antigen-specific hybridomas encapsulation could be omitted without appreciable cross-talk because target antibody signal was relatively infrequent. Finally, after allowing accumulation of sufficient antibody signal, samples were washed a final time and labeled with 0.75 µg/mL of hen egg lysozyme (Aviva Systems Biology) tagged with Alexa Fluor™ 647 (Microscale Protein Labelling Kit, Thermo Fisher) (HEL-647). All samples were sorted on a Sony SH800 flow sorter in single-cell mode with minor adjustments to the drop delay to account for the larger size of the particle nanovials (*44*).

### B cell secretion assay

Nanovials with 35 or 55 µm diameters were labeled with biotinylated anti-mouse CD45 and biotinylated goat anti-mouse IgG H&L chain antibodies (40 µg/mL and 60 µg/mL respectively) for 30 min. After incubation, nanovials were washed and resuspended in EasySep buffer. B cell lineage cells purified from splenocytes (see ‘Mouse immunization, serum isolation, and splenocyte isolation’) were seeded into a 24-well plate according to the section ‘Cell loading in nanovials’, and then incubated at 37°C for 1 hour to allow cells to adhere. To remove unattached cells from the background, samples were strained using a 37 µm reversible cell strainer and then particles were recovered by flipping the cell strainer and washing with washing buffer. After recovery, nanovial samples were concentrated by centrifuging particles and associated cells at 300 g for 3 min, aspirating, and then resuspending in 1 mL EasySep buffer. Samples were compartmentalized by pipetting with oil and surfactant (‘Dropicle formation and characterization’). Samples were then incubated for 2 hours to allow cells to secrete and to facilitate capture of secreted antibodies onto the associated nanovial matrix via goat anti-mouse IgG H&L sites. After the incubation period, nanovials and associated cells were transferred back into EasySep buffer by breaking the emulsions (see ‘Dropicle emulsion breaking’). Nanovial samples were then washed, and captured secretions were stained with AlexaFluor™ 647 conjugated ovalbumin (Invitrogen O34784) at a final concentration of 50 µg/mL. After 30 min of staining, samples were washed three times in a large volume of EasySep buffer and optionally stained with propidium iodide. An aliquot of the sample was then either sorted using a Sony SH800 FACS system based on high AF647 signal (e.g. the gate shown in Figure 6) or imaged with a fluorescence microscope in both brightfield and fluorescence channels in a well plate. Sorted nanovials were also imaged by fluorescence microscopy.

### Serum measurements, LOD and dynamic range experiments

For assessment of the dynamic range of the on-nanovial immunoassay for antibody capture and antigen-specific detection, 2 µL aliquots of nanovials were used for each condition. Nanovials with 55 µm diameters were labeled with biotinylated anti-mouse CD45 and biotinylated goat anti-mouse IgG H&L chain antibodies (40 µg/mL and 60 µg/mL respectively) for 30 min. After incubation, nanovials were washed and resuspended in washing buffer. Mouse serum (see ‘Mouse immunization, serum isolation, and splenocyte isolation’) at various dilutions was added to the particles and incubated for 2 hours. After the incubation period, particles were washed, and captured secretions were stained with AlexaFluor™647 conjugated ovalbumin (Invitrogen O34784) at a final concentration of 50 µg/mL. After 30 min of staining, samples were washed five times with washing buffer. Finally, particles were analyzed by fluorescence microscopy in both brightfield and fluorescence channels in a well plate and mean fluorescence determined for each serum dilution.

### Single-cell RT-PCR

Single HyHel-5 hybridoma cells or single nanovials loaded with HyHel-5 cells were separately sorted into 96-well plates (Sony SH800) containing 5 mM DTT, 0.7% NP-40, 5 µM random hexamers, and 1U RNasin Plus in PBS and stored at -80°C until analysis. For first-strand cDNA synthesis, sorted cells were thawed on ice, incubated at 65 °C for 1 min, then incubated on ice for 2 min. Each well then received 10 µL of 2X RT Buffer, 5 mM MgCl_2_ and 0.01 M DTT containing 40 U RNasin Plus and 200 U SuperScript III Reverse Transcriptase (Thermo Fisher). The reaction was incubated at 42 °C for 10 min, 25 °C for 10 min, 50 °C for 1 hour, then 94 °C for 5 min. The cDNA product from each single cell was then amplified for heavy chain in a sequence of nested RT-PCR reactions using primers optimized by von Boehmer et al (*45*). All PCR reactions contained 1X HF Buffer, 200 µM dNTPs, and 0.02 U/µL Phusion High Fidelity DNA Polymerase (Thermo Fisher) in a final reaction volume of 20 µL. The initial, preamplification RT-PCRs contained a mixture of forward primers at a final concentration of 0.3 µM, the reverse primer at a concentration of 0.2 µM, and 2 µL of the reverse-transcribed cDNA. The thermocycler conditions included an initial denaturation at 98 °C for 5 min followed by 50 cycles of 30 sec at 98 °C, 30 sec at 46 °C, and 30 sec at 72°C then a final extension at 72 °C for 10 min. The second, nested RT-PCRs contained 0.2 µM each of the forward and reverse primers and 2 µL of the preamplification RT-PCR product. The thermocycler conditions were the same as in the preamplification PCR, but with a 57 °C annealing temperature. The second PCR products were visualized on a 1.5% agarose gel with SYBR Safe DNA gel stain (Thermo Fisher).

## Supporting information

Video S1 Nanovial Assay Workflow

Video S2 Nanovial Settling

## Acknowledgements

We acknowledge support from the National Institutes of Health Grants R21GM126414, T32GM008042, T32AR071307, R01EB029455, R01EY031097, U01CA232563, a Maryland Stem Cell Research Fund Discovery Award, and the Simons Foundation Math+X Investigator Award #510776. Device mold fabrication was completed using equipment provided by the Integrated Systems Nanofabrication Cleanroom at the California NanoSystems Institute at the University of California, Los Angeles. Sorting experiments were performed in the UCLA Jonsson Comprehensive Cancer Center (JCCC) and Center for AIDS Research Flow Cytometry Core Facility that is supported by National Institutes of Health awards P30CA016042 and P30AI028697, and by the JCCC, the UCLA AIDS Institute, the David Geffen School of Medicine at UCLA, the UCLA Chancellor’s Office, and the UCLA Vice Chancellor’s Office of Research. We thank Richard Willson for kindly providing the HyHel-5 hybridoma line for these studies. The authors acknowledge Philip Scumpia for assistance with mouse immunization studies. The authors acknowledge the help of Aiden Di Carlo in editing the secretion assay workflow video.

## Author Contributions

J.D., R.Di, and D.D. conceptualized the overall nanovial platform and designed experiments throughout the entire work. R.Da. contributed to designing and interpreting flow cytometry experiments and single-cell RT PCR experiments and B cell secretion experiments. J.D. designed the initial fabrication workflow and performed characterization experiments. J.D., R.Di, S.Z., S.L., S.U., and D.K. contributed to production and optimization of nanovials for the experiments. J.D, R.Di., M.A, J.E. M.Z., D.K. S.U., and P.K. performed cell loading and cell binding experiments. J.D., R.Di., and S.Z. performed and analyzed nanovial emulsification experiments. J.D., R.Di., and M.V. developed the CHO cell secretion protocols and performed characterization experiments. P.J.K. and M.K. performed HEK293 transfection studies which J.B.S. supervised. R.Di., M.A., S.L., J.E, and S.Z., performed hybridoma and B cell secretion assay experiments. M.A. and J.E. performed mouse immunizations and harvested cells for B cell studies. M.P.M. and A.C.S. performed single-cell RT-PCR experiments. K.H. and A.L.B. developed the framework for calculating the volume energy curves and volume splitting curves. J.D., R.Di, and D.D. wrote the manuscript with input from all authors. J.D. prepared final figures with input from all authors.

