## Supplemental Information for "Massively parallel encapsulation of single cells with structured microparticles and secretion-based flow sorting"

### Supplementary Information

Massively parallel encapsulation of single cells with structured microparticles and secretion-based flow sorting

**Video S1. Protocol video demonstrating the complete secretion assay workflow step by step.**

**Video S2. Time-lapse of a suspension of nanovials being seeded into a well plate.** Settling takes less than 5 minutes and a large fraction of particles settle with their exposed cavities oriented upright enabling cell loading.

**Figure S1. Nanovial fabrication using an aqueous two-phase system combined with droplet microfluidics.**

**Figure S2. Nanovial morphology can be tuned by adjusting the concentrations of PEG and dextran in the droplets.**

**Figure S3. Compatibility of a range of nanovial sizes with different cell types and instruments.**

**Figure S4. Time-lapse images of nanovials seeded into a well plate.**

**Figure S5. Energy minimalization theory predicts monodisperse dropicle formation.**

**Figure S6. Characterization of cell viability and growth after dropicle formation and release.**

**Figure S7. Detailed overview of high IgG producer enrichment workflow.**

**Figure S8. Analysis of single-cell IgG secretion distributions for transiently transfected cells on nanovials.**

**Figure S9. Detailed overview of hybridoma and B cell secretion assay.**

**Figure S10. Single-cell RT-PCR gel results for free hybridomas and hybridomas loaded on nanovials.**

**Figure S11: Assessment of the dynamic range of nanovials for capture and antigen-specific detection of antibodies.**

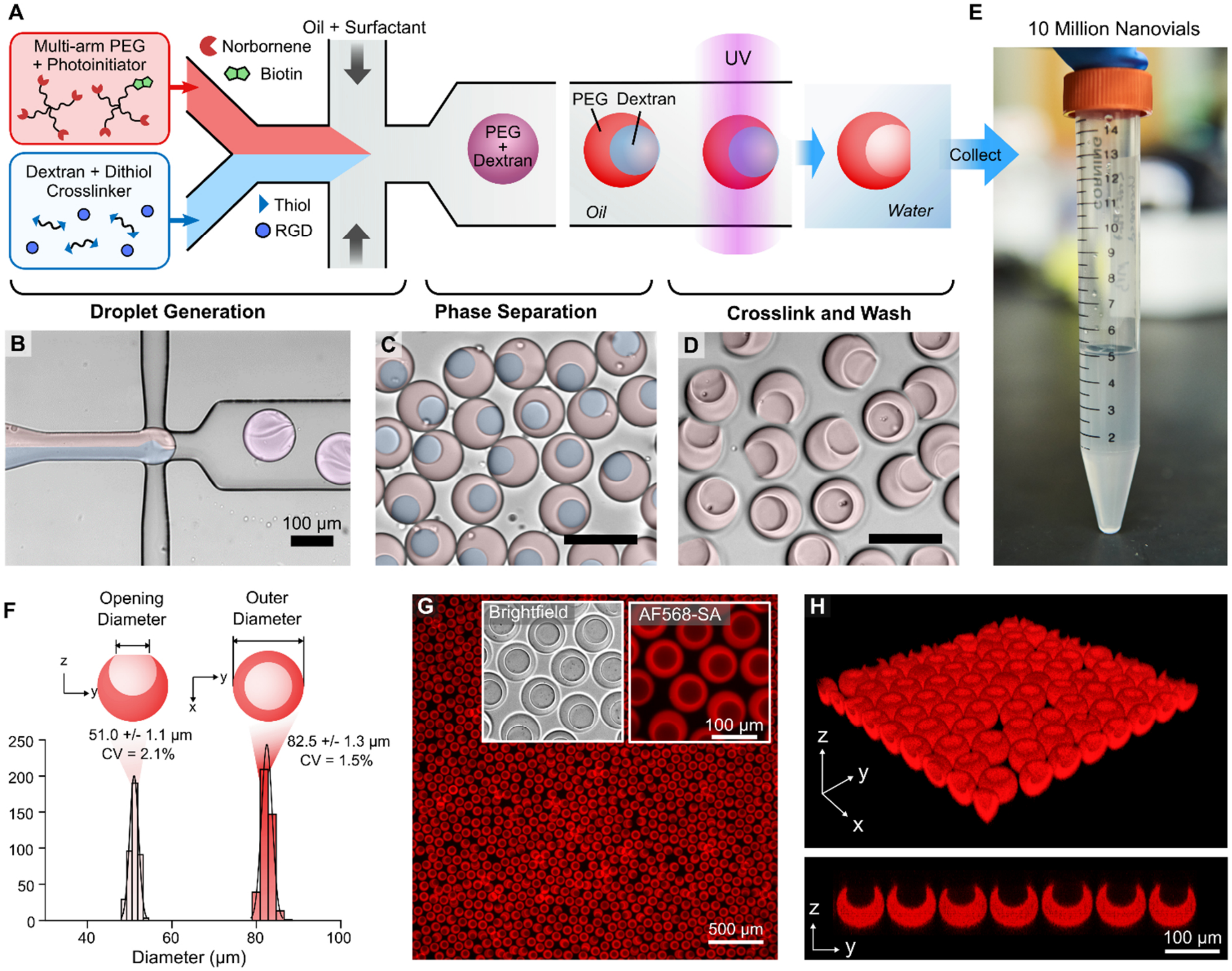


**Figure S1. Nanovial fabrication using an aqueous two-phase system combined with droplet microfluidics.** **(A)** A solution comprised of UV reactive PEG and photoinitiator is co-flowed with a solution containing dextran, dithiol crosslinkers, and RGD peptides in a microfluidic droplet generator. **(B)** A third solution of oil and surfactant is injected into the device to generate water-in-oil droplets at a rate of ~1000 Hz. **(C)** After droplet formation, the PEG and dextran undergo phase separation resulting in two distinct regions in the droplet. The droplets are exposed to UV light at the end of the device to crosslink the PEG-rich portion of the droplet, while the dextran-rich region remains as a liquid. **(D)** The resulting microparticles are then collected, washed to remove oil and dextran, and stored for later use. **(E)** Photograph of a 15 mL conical tube with 10 million particles fabricated in ~3 hours. **(F)** Nanovials fabricated using this approach are highly monodisperse with an outer diameter CV of 1.5% and a cavity opening diameter CV of 2.1% (n = 409). Particle uniformity was calculated by analyzing fluorescence microscopy images using a custom image analysis algorithm in MATLAB. **(G)** Fluorescence microscopy image of biotinylated nanovials stained with Alexa Fluor$^{TM}$ 568 Streptavidin (AF568-SA). A large fraction of the seeded particles settles with their cavities exposed upright. **(H)** Nanovial cavity morphology and upright orientation was confirmed using confocal microscopy. Microscopy images in B, C, and D have overlayed color to aid in visualization.

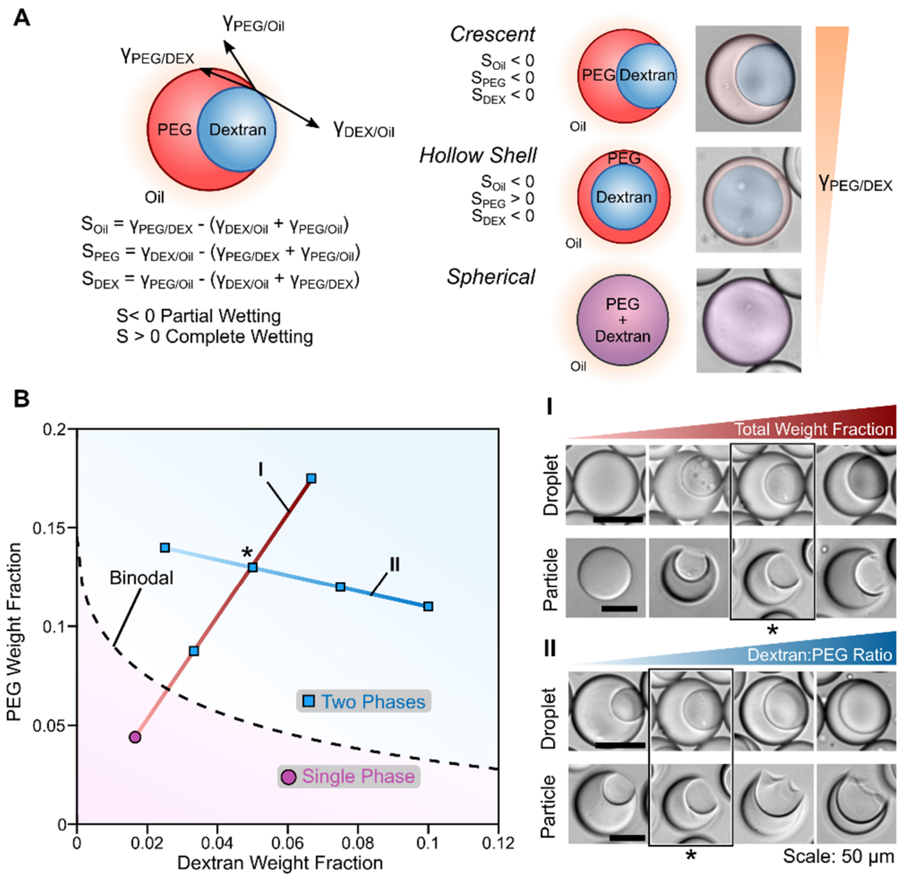


**Figure S2. Nanovial morphology can be tuned by adjusting the concentrations of PEG and dextran in the droplets.** (**A**) The morphology of nanovials is dictated by a balance of interfacial tensions ($\gamma$) between the different phases. $\gamma_{PEG/Oil}$ and $\gamma_{DEX/oil}$ are expected to vary minimally, the balance of interfacial tensions and the resulting morphology is expected to be dictated mostly by a change in $\gamma_{PEG/DEX}$. (**B**) Experimentally determined phase diagram of 4-arm PEG norbornene, and dextran. At very low concentrations phase separation does not occur resulting in a spherical particle (**I**). As the total concentration of PEG and dextran is increased above the binodal line, phase separation occurs enabling fabrication of cavity-containing microparticles. (**I**) The relative opening diameter of the particle cavity is increased by increasing the total polymer concentration and thus $\gamma_{PEG/DEX}$ resulting in a relative opening size of 48 – 63% of the outer diameter. (**II**) By adjusting the concentration ratio of dextran and PEG, particles can be fabricated with different relative cavity sizes. Here the relative cavity diameter is from 34 – 76% the outer diameter. Particles fabricated using a PEG concentration of 13% and dextran concentration of 5% (denoted by *) were found to have high structural integrity while maintaining a relatively large cavity opening to enabled efficient cell loading. This condition was used for all other experiments in this work.

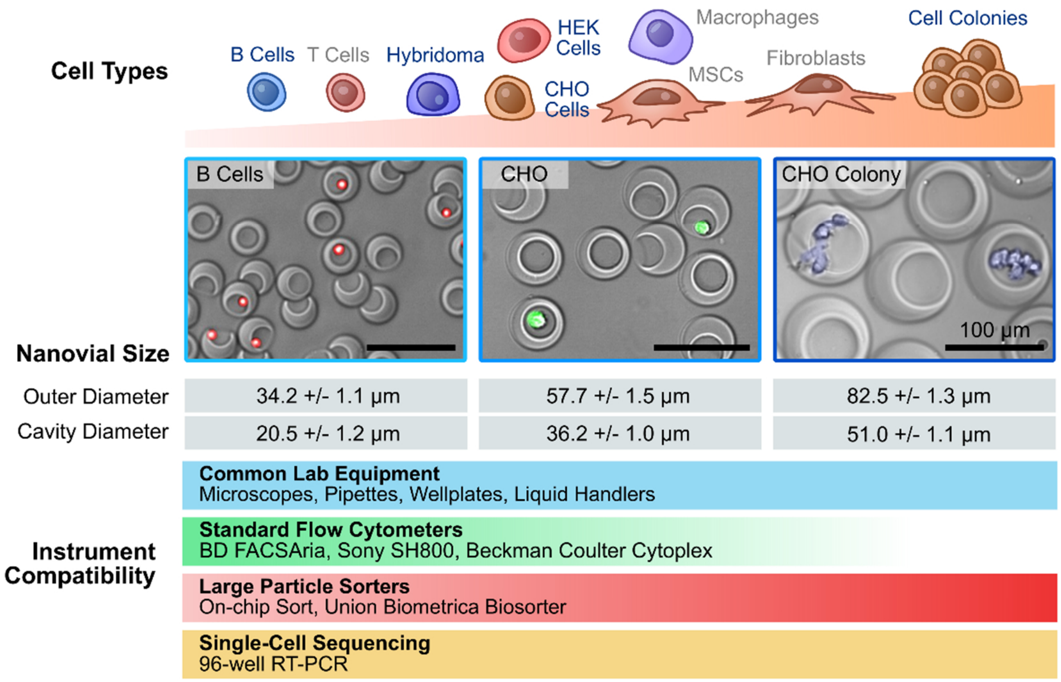


**Figure S3. Compatibility of a range of nanovial sizes with different cell types and instruments.** Nanovials with high uniformity were fabricated with mean outer diameters ranging from 35 to 82 µm and respective mean cavity diameters of 20 to 51 µm. Smaller nanovials are more compatible with standard flow cytometers which are more accessible and allow higher throughputs, but are limited to smaller cell types. Larger nanovials are compatible with larger cells and cell colonies/clusters containing more cells, but are compatible with fewer commercially available flow sorters. Notably all sizes are compatible with microscopy, pipettes and liquid handlers. Cell labels highlighted blue identify cells that were shown to be compatible with nanovials in this work. CHO colonies are false labeled blue to aid in visualization. B cell and CHO cell microscopy images are overlayed brightfield and fluorescence images. Scale = 100 µm.

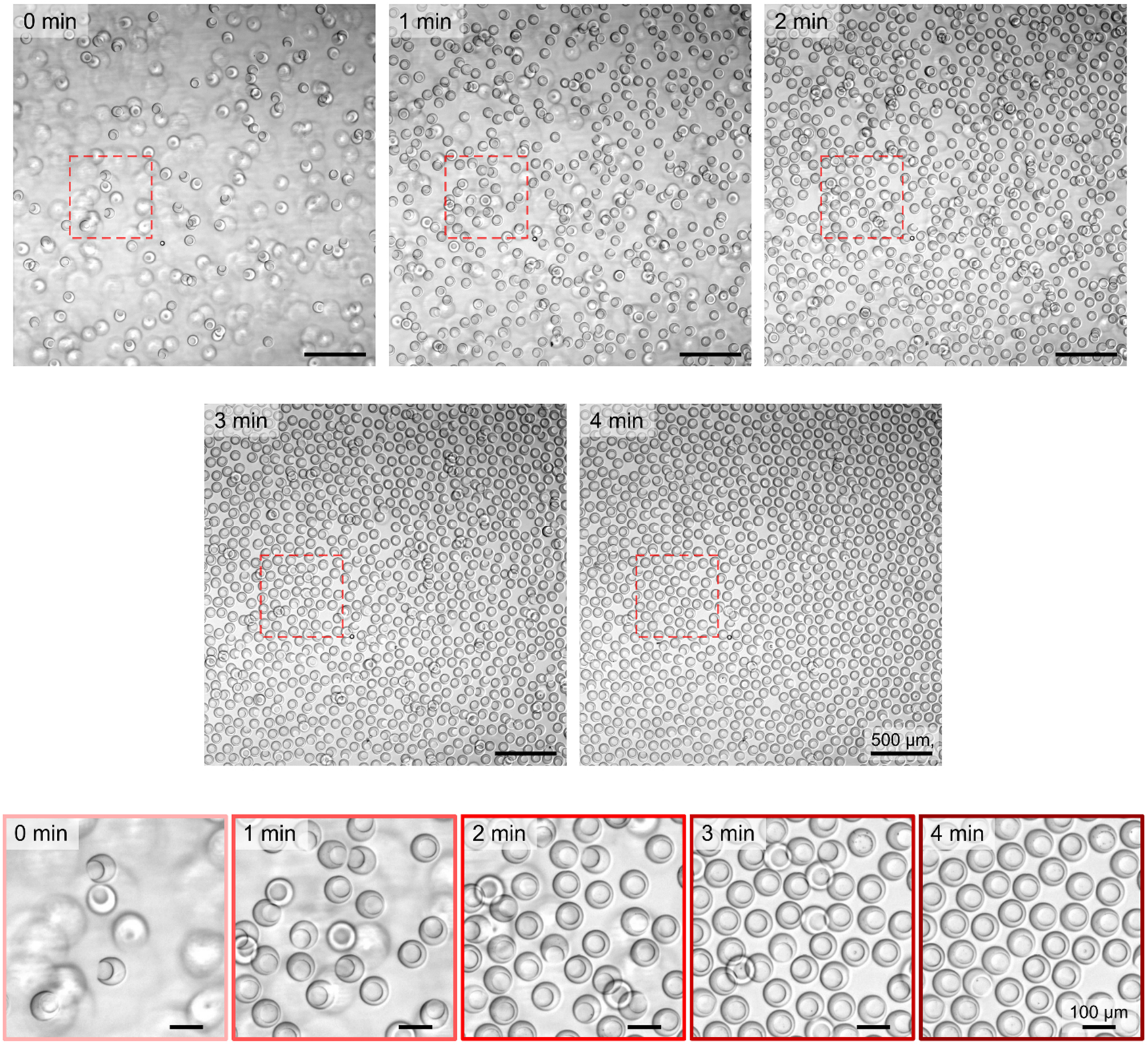


**Figure S4. Time-lapse images of nanovials seeded into a well plate.** To prepare nanovials for cell seeding they are pipetted into a well plate by dispensing liquid up and down in circular motions. Due to the unique morphology of the nanovials they tend to settle with their cavities oriented upright and displace adjacent nanovials aside to form a monolayer.

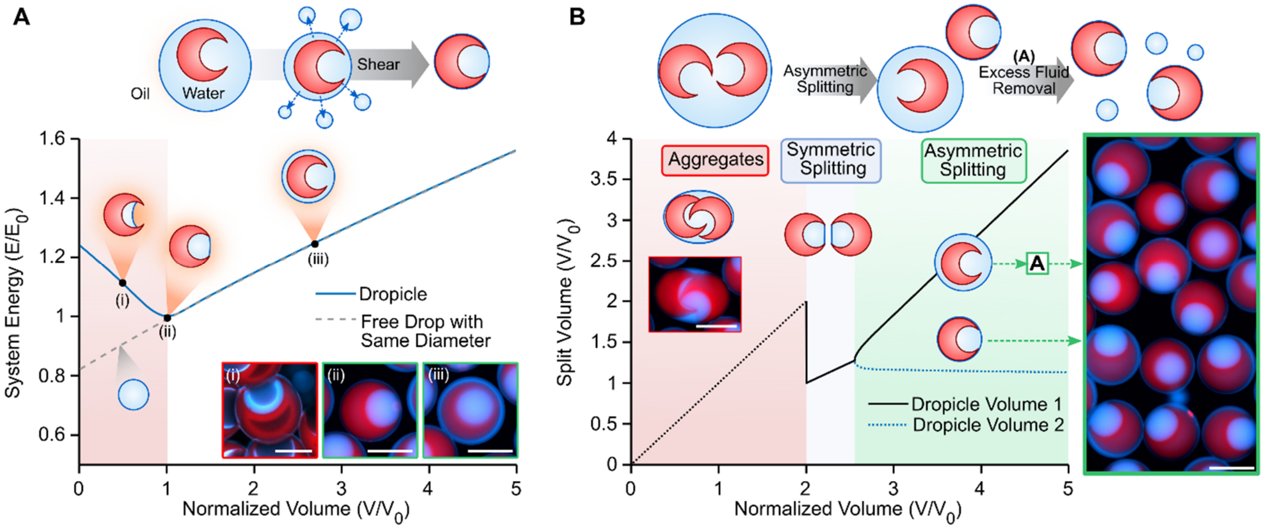


**Figure S5. Energy minimization theory predicts monodisperse dropicle formation.** **(A)** Simulated volume-energy (V-E) curves show that droplets formed with nanovials (dropicles) have a minimal energy configuration when the size of the droplet surrounding it approaches the size of the nanovial. Here the system energy and volume are normalized by the minimum energy values (E0 and V0, respectively). A free droplet of the same outer diameter follows along the same energy curve until the minimal energy point. As energy is added to the system (e.g. by shearing with a pipette) excess fluid is shed from the nanovial (**iii**), until the minimal energy volume is reached (**ii**). Lower volumes could only be achieved by drying out excess fluid via evaporation (**i**). **(B)** On the basis of the V-E curve in (**A**), splitting of a drop containing two nanovials is theoretically expected to depend on the total fluid volume. At volumes below twice $V_0$ nanovials are expected to remain as aggregates. At slightly larger volumes if enough energy is added the volume is expected to split evenly based on energy minimization theory. At larger volumes the fluid is expected to distribute asymmetrically between nanovials; one with a smaller volume approaching $V_0$ and one with the remainder of the fluid. The nanovial with the larger volume may undergo additional volume reduction via shearing until reaching $V_0$. Scale: 50 µm.

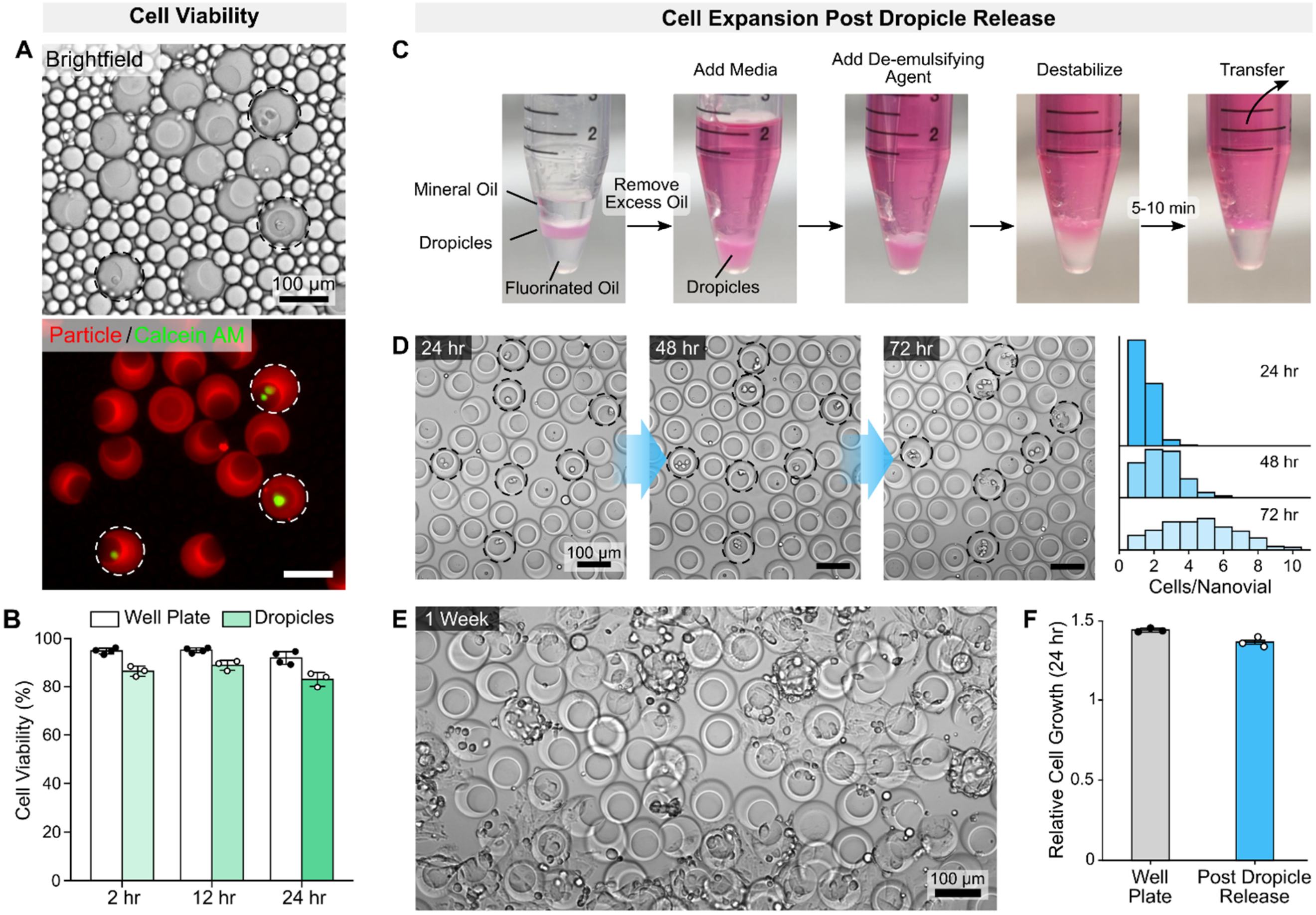


**Figure S6. Characterization of cell viability and growth after dropicle formation and release.** (**A**) Brightfield and fluorescence microscopy images of CHO cells encapsulated in dropicles. Biotinylated nanovials are stained with Alexa Fluor$^{TM}$ 568 streptavidin and cells are stained with calcein AM. (**B**) Cells maintained high viability after dropicle formation and release. Viability was assessed by staining with calcein AM and propidium iodide after recovering nanovials and cells from droplets. A live/dead assay was performed with a minimum of n = 3 samples, cell number > 1000 per condition. (**C**) Process for recovering nanovials and cells from emulsions. (**D**) Cells initially remain in the nanovial cavities after releasing them. Colonies derived from single cells remain in the particle cavities during initial expansion. (**E**) After significant accumulation of cells in the cavities, cells begin to spread to the outside of the cavities and onto the well plate surface. (**F**) Cell growth in nanovials post dropicle release is comparable to cell growth on standard well plates (control) (n = 3). Growth is characterized by fold change in cell number over a 24 hr period as characterized by counting cell number using CellTracker$^{TM}$ staining.

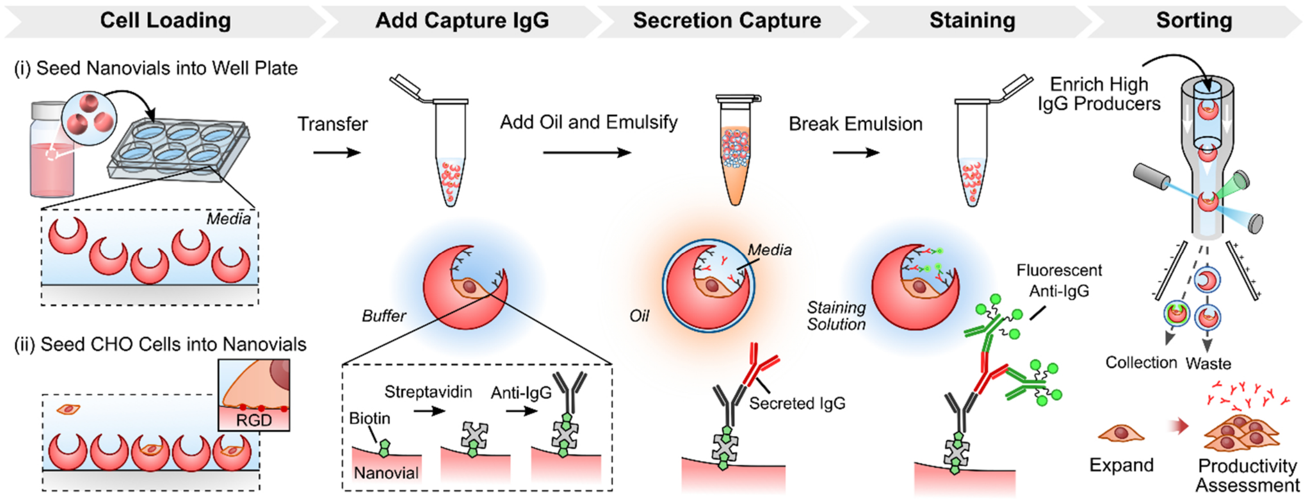


**Figure S7. Detailed overview of high IgG producer enrichment workflow.** RGD coated nanovials are first seeded into a well plate and settle with cavities exposed upright. CHO cells are then seeded into the exposed cavities and incubated to allow attachment via integrin binding. Nanovials and attached cells are recovered by pipetting and transferred into a centrifuge or conical tube. Nanovials are then modified sequentially with streptavidin and biotinylated anti-IgG antibodies. The nanovials are then emulsified to prevent crosstalk and incubated in a CO2 incubator such that secreted antibodies from cells accumulate. Nanovials, cells, and associated secretions are transferred back into a water phase, labeled with fluorescent secondaries, and analyzed with flow to measure amount of secreted product. The higher producing cells, as measured by fluorescence, are sorted out and expanded in a well plate for downstream productivity assessment. The full secretion assay workflow is completed in a single day and cells are expanded for a week after enrichment for downstream assessment via bulk well plate ELISA.

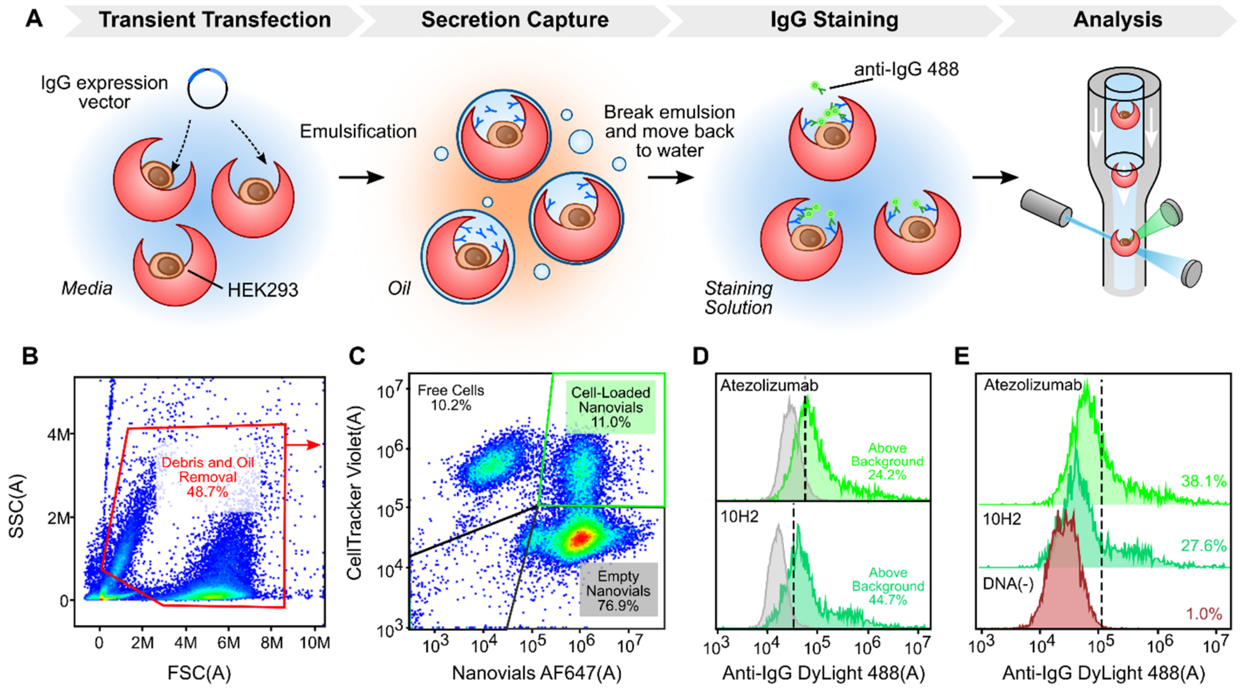


**Figure S8. Analysis of single-cell IgG secretion distributions for transiently transfected cells on nanovials. (A)** Biotinylated HEK293 cells stained with CellTracker™ Violet were loaded onto streptavidin coated nanovials. After cell binding, biotinylated anti-IgG was bound to remaining streptavidin sites. Cells are then transiently transfected with an IgG expression vector and incubated for 16 hrs. Cells and nanovials were then emulsified in oil and incubated for an addition 32 hrs to accumulate secretion signal. Cells and nanovials were transferred back to buffer solution and stained with fluorescent secondary antibodies against human IgG and analyzed using flow cytometry. **(B)** Representative forward and side scatter gating strategy is shown. Debris and residual oil are removed by excluding low scatter signal. **(C)** To accurately differentiate subpopulations, cells were tagged and gated based on CellTracker™ Violet signal and nanovials were tagged and gated based on AlexaFluor™ 647 signal. **(D)** A fraction of cell-containing nanovials were observed to have IgG signal above empty nanovials indicating cell secretion-specific signal. **(E)** For both constructs we see signal above a negative DNA control sample indicating cell secretion-specific signal.

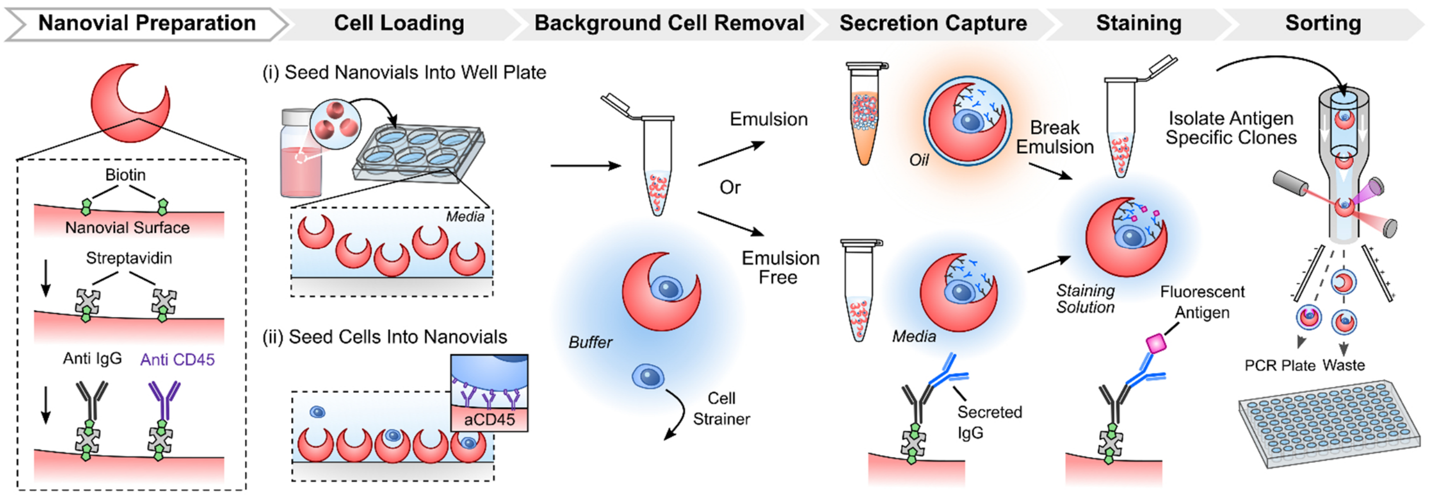


**Figure S9. Detailed overview of hybridoma and B cell secretion assay.** Biotinylated nanovials are prepared prior to the assay by coating with streptavidin followed by labeling of the nanovial surface with biotinylated anti IgG for secretion capture and biotinylated anti CD45 for cell binding. Nanovials are seeded into a well plate and hybridomas or B cells are then loaded into the exposed cavities. Cells are incubated to facilitate binding via CD45 surface markers. Samples are recovered by pipetting and unbound cells are removed using a reversible cell strainer. Samples are then either emulsified or left in media and allowed to incubate to accumulate secreted IgG on the nanovial surface. Captured secretions are then labeled with fluorescently tagged antigens, washed, and analyzed and sorted using a flow sorter.

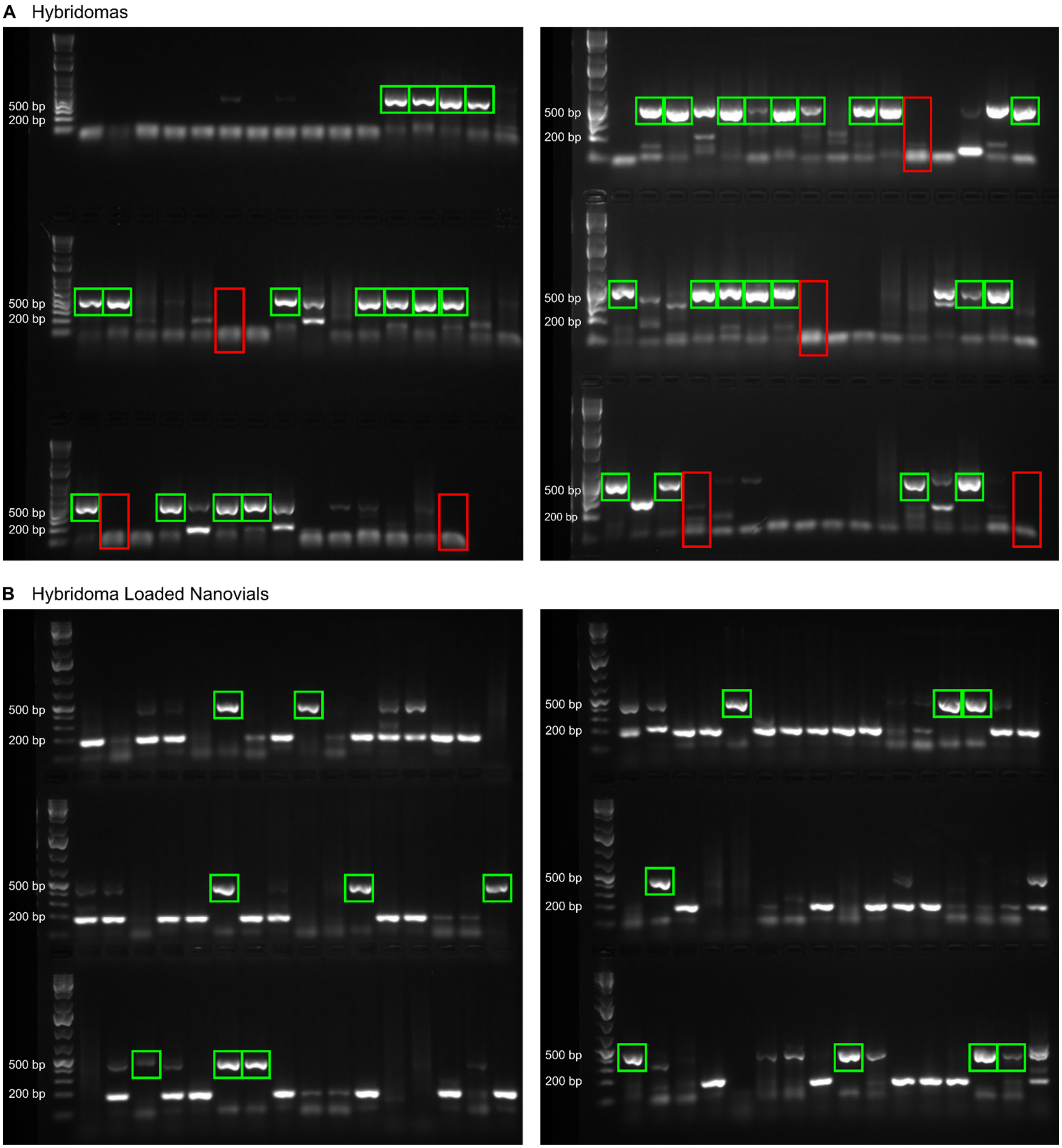


**Figure S10. Single-cell RT-PCR gel results for free hybridomas and hybridomas loaded on nanovials.** RT-PCR amplified heavy chain DNA products were visualized on agarose gels from single wells of a 96 well plate following FACS sorting of single hybridoma cells (A) and single hybridoma cells loaded on nanovials (B). (**A**) Properly amplified heavy chain products were detected in 35/87 single-cell sorted wells (green boxes). 0/7 negative control wells yielded amplified products (red rectangles). Based on preliminary cell sorts, 81% of wells are expected to contain a cell, resulting in an amplification efficiency of 50%. (B) Properly amplified heavy chain products were detected in 16/96 single-cell sorted wells (green boxes). Based on preliminary nanovial sorts 31% of wells are expected to contain a cell-loaded nanovial resulting in an amplification efficiency of 54%.

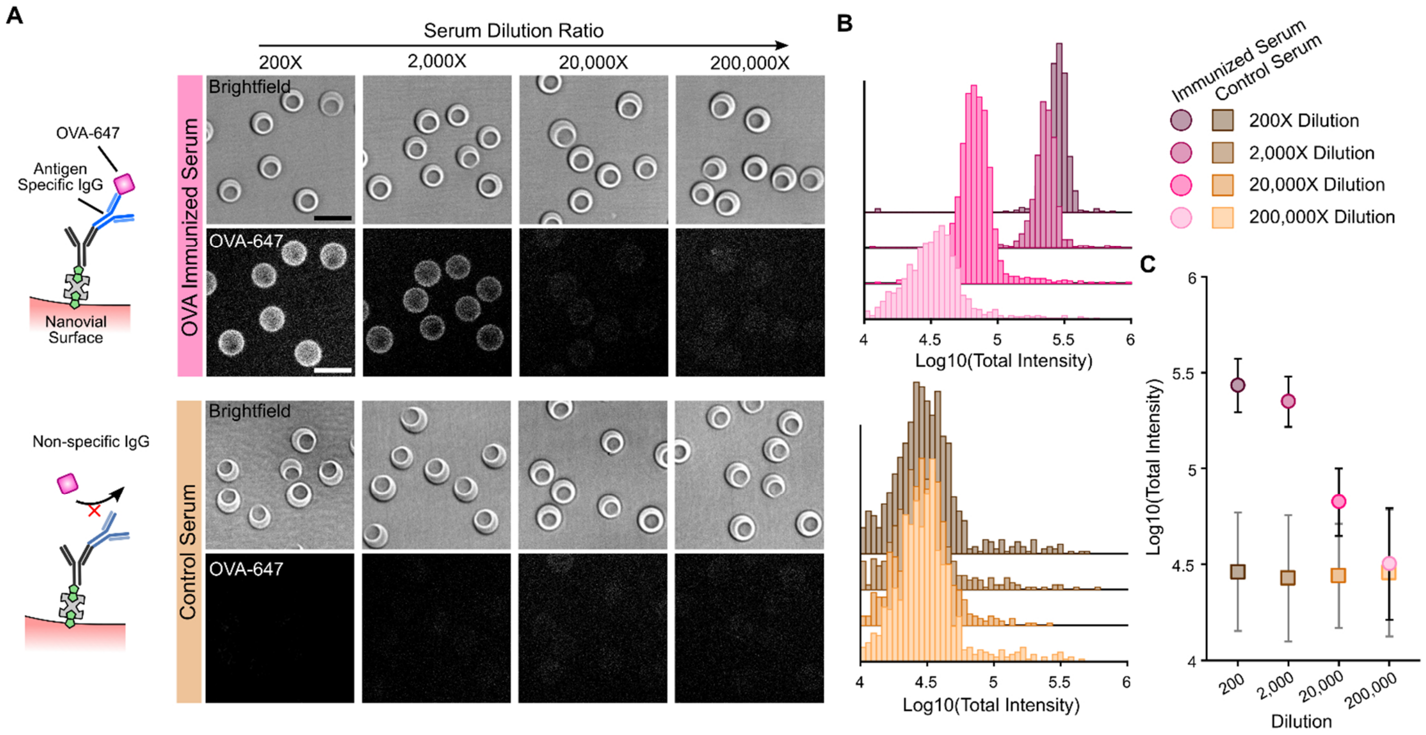


**Figure S11**: **Assessment of the dynamic range of nanovials for capture and antigen-specific detection of antibodies**. (**A**) Biotinylated anti-Mouse IgG H+L antibody-coated nanovials were incubated with serial dilutions of sera from ovalbumin (OVA)-immunized and non-immunized mice and stained with AlexaFluor$^{TM}$ 647 conjugated OVA. (**B**) Nanovials were detected with antigen-specific antibodies in immunized mice sera at dilutions down to 1:20,000 and show no non-specific binding in control serum using fluorescence microscopy. (**C**) Data is summarized as a function of serum dilution, showing the functionality of the on-nanovial immunoassay for OVA-specific IgG.
